# Unexpected variability of allelic imbalance estimates from RNA sequencing

**DOI:** 10.1101/2020.02.18.948323

**Authors:** Asia Mendelevich, Svetlana Vinogradova, Saumya Gupta, Andrey A. Mironov, Shamil Sunyaev, Alexander A. Gimelbrant

## Abstract

RNA sequencing and other experimental methods that produce large amounts of data are increasingly dominant in molecular biology. However, the noise properties of these techniques have not been fully understood. We assessed the reproducibility of allele-specific expression measurements by conducting replicate sequencing experiments from the same RNA sample. Surprisingly, variation in the estimates of allelic imbalance (AI) between technical replicates was up to 7-fold higher than expected from commonly applied noise models. We show that AI overdispersion varies substantially between replicates and between experimental series, appears to arise during the construction of sequencing libraries, and can be measured by comparing technical replicates. We demonstrate that compensation for AI overdispersion greatly reduces technical variation and enables reliable differential analysis of allele-specific expression across samples and across experiments. Conversely, not taking AI overdispersion into account can lead to a substantial number of false positives in analysis of allele-specific gene expression

---

RNA sequencing (RNA-seq) is a widely used technology for measuring RNA abundance across the whole transcriptome ^1^. An especially informative approach to RNA-seq analysis in samples from humans and other diploid organisms is comparison of the activity of the parental alleles. Allele-specific analysis of gene expression can reveal epigenetic gene regulation associated with imprinting ^2^, X-chromosome inactivation ^3^, and autosomal monoallelic expression ^4–8^. The maternal and paternal copies of a gene share the same cell nucleus and therefore are both influenced by the rest of the genome in the same way. Consequently, allelic imbalance (AI) in expression can be highly sensitive to cis-regulatory mechanisms ^9,10^. Accordingly, AI analysis has been used to uncover gene regulatory effects in a growing number of studies ^11–14^.

Accurate estimation of AI is thus important for quantitative understanding of genetic and epigenetic mechanisms of gene regulation. Efforts to increase AI estimation accuracy have mostly focused on the data analysis ^15–20^, based on the assumption that consistency in measuring total RNA abundance translates to accurate measurement of each of the alleles separately. Here, we show that this implicit assumption is incorrect.

Based on analysis of newly generated datasets and publicly available data, we reveal a previously neglected, major source of technical variation in AI measurements in RNA-seq experiments. This indicates that analyses based on a single RNA-seq replicate and aiming to identify genes with significant AI can result in an unexpectedly large fraction of false positives. To correct for this issue, we devised a specific measure of AI overdispersion, Quality Correction Constant (QCC), derived from comparison of replicate RNA-seq libraries. QCC can be used to control for this unexpected source of variation, and its use results in more accurate AI estimates. Finally, we outline several use cases for this improved approach, including differential analysis of allele-specific expression.

## Results

### Datasets used

There are two principal variables involved in generation of RNA sequencing libraries: (a) different protocols can be used; and (b) with a given protocol, library preparation can start with different amounts of RNA. To probe both variables in a compact way, we generated three sets of poly-A enriched RNA-seq libraries from the same RNA. Each set (“experiment”) consisted of six libraries prepared in parallel starting from the same biological sample: total RNA extracted from the kidney of a female mouse (129S1×Cast/Ei F1 cross, with genome-wide SNP density of ~1/118 bp). Libraries for experiment 1 (“NEBNext (100ng)”) were prepared using a protocol for large amounts of input RNA (100 ng of total RNA, see details in Methods). Experiments 2 and 3 featured libraries prepared using SMART-Seq v4 Ultra Low Input RNA Kit (Clontech) with amounts of input total RNA bracketing the recommended range - 10 ng and 100 pg, respectively (“SMARTseq (10ng)” and “SMARTseq (0.1ng)”).

Properties of the libraries and the sequencing data are summarized in **Suppl. Table S1,** together with the properties of the previously published, publicly available datasets we analyzed, including data from mouse ^21^ and human ^22,23^ (**Suppl. Tables S2** and **S3**).

### Replicate RNA-seq libraries exhibit unexpected level of variation in the AI estimates

In the analysis of allele-specific expression from sequencing data, the use of a single RNA-seq library is a common practice ^16,24^. Indeed, comparison of (non allele-specific) RNA abundance values between two replicate RNA-seq libraries shows good agreement, even only counting the reads that cover SNPs (**Fig.1a**). There appears to be much less agreement when comparing AI values calculated from the same two replicate libraries (**Fig.1b**). This could be expected, since AI values are proportions which amplify small variations.

**Figure 1.**
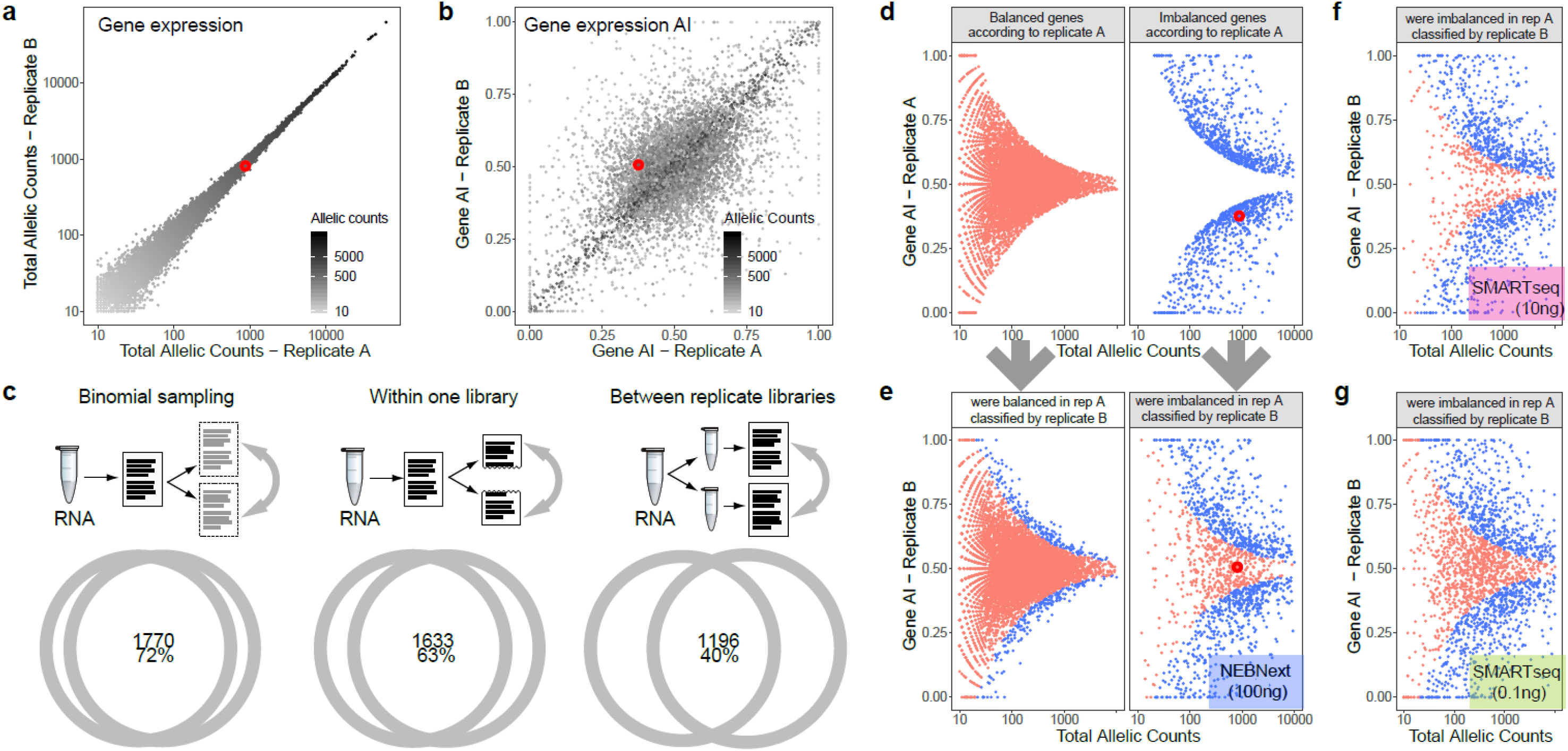
Allelic imbalance estimates disagree between technical replicates, and that disagreement varies between RNA-seq experiments. **a, b:** Comparison of two replicate RNA-seq libraries prepared from the same RNA [NEBNext(100ng) experiment]. Total allelic counts reflected in greyscale; only genes with total allelic counts >10 are shown. *Red circle* highlights an example gene (*Stolm2*, in top 15% by expression). RNA from 129×CastF1 mouse kidney. **a**: Comparison of AI values [maternal allelic counts/(total allelic counts)]; **b**: comparison of total allelic counts. **c:** Concordance of “allelically imbalanced” genes (*H*_*0*_ of AI=0.5 is rejected by binomial test; p=0.05 with Bonferroni correction) between two sets of 25M RNA-seq reads [SMARTseq(0.1ng)]: *Left*: sets of reads in a binomial relationship with each other (sets independently subsampled from all 52M reads from the same RNA-seq library); *Middle*: sets obtained by splitting in half 50M reads from the same RNA-seq library; *Right*, sets sampled from two replicate libraries prepared in parallel. **d-g:** Comparison of gene classification between two technical replicates. <u>Red</u>: *H*_*0*_ of AI=0.5 is **not** rejected by binomial test; p=0.05 with Bonferroni correction; <u>Blue</u>: rejected by the same test. 30M reads were sampled per each replicate. **d, e:** same data as (a) and (b). Example gene (*Stolm2*) highlighted. **d**: genes classified based on data from Replicate 1; **e**: genes grouped as in (d) but classified and colored based on data from Replicate 2. **f, g:** analysis as in right panel in (e). **f**: SMARTseq(10ng) experiment; **g**: SMARTseq(0.1ng) experiment.

To understand how this increase in disagreement affects the reproducibility of AI measurements, we used a simple, commonly used procedure ^25,26^: whether null hypothesis of balanced gene expression (H_0_ of AI=0.5) is rejected by the binomial test (p=0.05 with Bonferroni correction). To establish the baseline, we first compared two sets of reads in a binomial relationship with each other (**Fig.1c**, *left*; two sets independently subsampled from all reads from the same RNA-seq library); in this comparison, 1,770 genes were concordant between the two sets (72% of the number of genes imbalanced in at least one of the sets). Two sets of reads sampled without replacement from the same RNA-seq library showed lower concordance (**Fig.1c**, *middle*), which can be explained by overdispersion known to be present in RNA abundance analyses of RNA-seq experiments ^27^. It is commonly thought that technical overdispersion can be sufficiently accounted for by within-replicate analysis (approaches include comparing sets of reads sampled without replacement from the same read pool ^24^ or bootstrapping ^28^). However, the concordance between a pair of technical replicates was much lower still (**Fig.1c**, *right*), indicating presence of additional noise, not detectable by analysis within a single library.

We then assessed the distribution of AI estimates for genes whose classification was different in two replicates (**Fig.1d,e**). Using AI estimates from one replicate, we divided genes into balanced and imbalanced (**Fig.1d**). Within these groups, we then used the AI values from another replicate to re-classify genes as (im)balanced. For the pair or replicates shown in **Fig.1d,e**, 5% of genes “balanced” in replicate A are classified as imbalanced in replicate B (*left*), and 30% of “imbalanced” genes in A are balanced in B (*right*).

Note that the observed AI values for genes with discordant binomial test calls between two replicates are not concentrated around the boundary determined by the binomial test (as an illustration, consider the highlighted gene in **Fig.1a,b,d,e**). This strongly suggests that a simple binomial noise assumption incorrectly describes the observed dispersion of AI values (see **Suppl. Note S1**).

Furthermore, the concordance rates were strikingly different when compared across different experiments (**Fig.1e, f, g**). At the same time, the concordance was similar for pairs of replicates within the same experiment: 49.2%±4.6 (s.d.) for experiment 1 (52.3±1.2 when one outlier replicate was removed), 61.8±0.6 for experiment 2 and 39.6±1.0 for experiment 3. We also analyzed several publicly available datasets ^21,22^ (**Suppl. Fig.S1**, **Suppl. Tables S2** and **S3**). When multiple replicates were available, we also observed that the concordance was similar for pairs of replicates within an experiment. We thus conclude that the AI overdispersion we observe is experiment-specific.

### Estimation of AI overdispersion from observed and modeled data

In order to quantify the experiment-specific overdispersion between a pair of replicate libraries, we assess how its experimentally observed value compares to the expected value from a fitted model. To discretize the model, the analysis is performed on genes binned by total allelic coverage (**Fig.2**).

**Figure 2.**
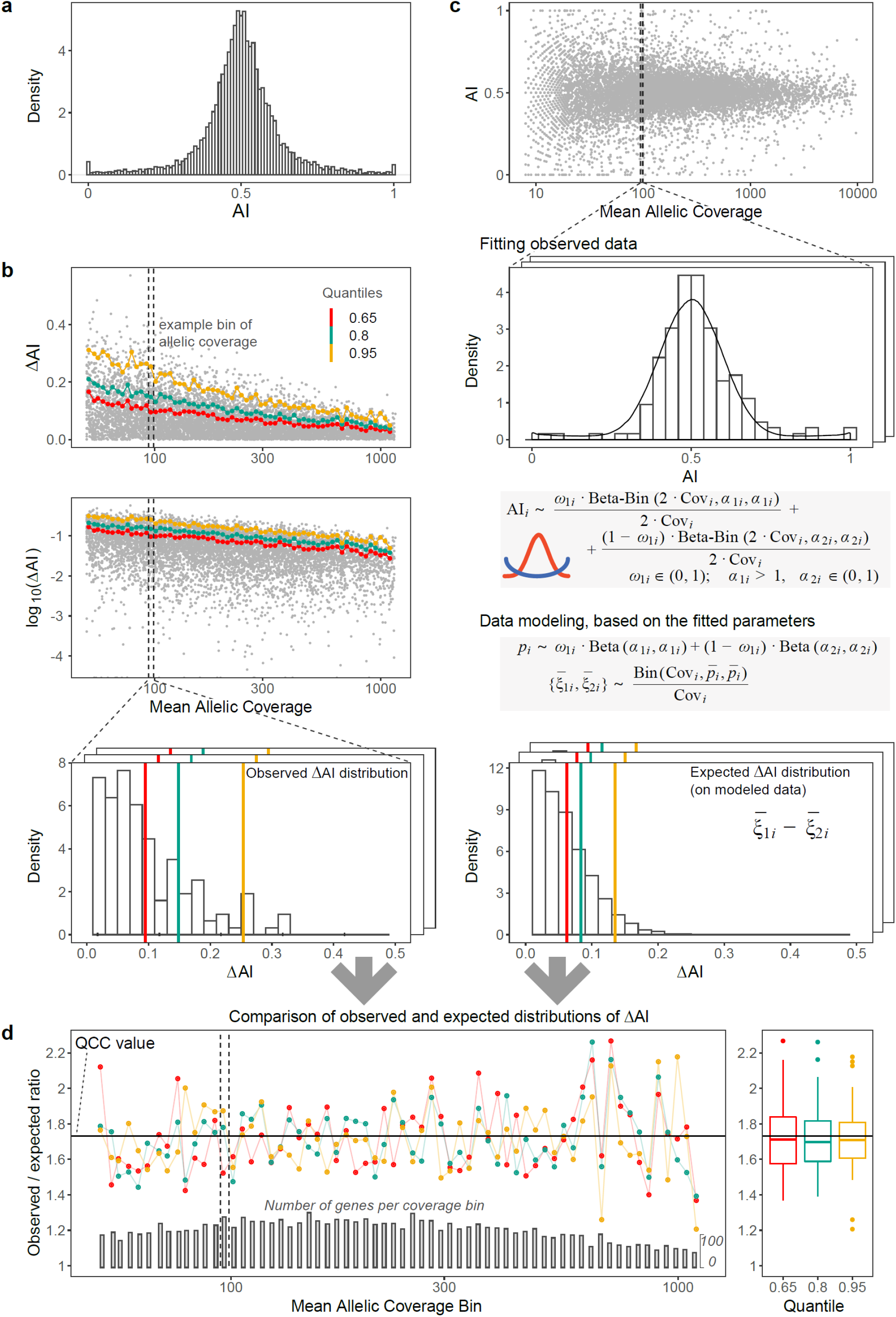
Derivation of Quality Correction Constant from observed and modeled AI differences between technical replicates. **a:** Distribution of AI for all genes in six pooled replicates (180M reads total) of SMARTseq (10ng) experiment. SNP coverage threshold of 10 or more reads was applied to the pooled data. **b:** Calculation of the observed distributions of AI differences between replicates. *Top*: After sampling equal number of reads from two technical replicates, AI is calculated for each gene. Plotted is the absolute difference of AI between replicates (**Δ**AI) against mean SNP coverage. Genes are binned by log coverage (an example bin is shown). For each bin, quantiles are calculated; shown are 65th (red), 80th (green), and 95th (orange) quantiles. *Middle*: same as top, in log-log coordinates. Note that quantiles are lying along straight lines in log-log scale. *Bottom*: Distribution of the observed **Δ**AI values in a selected bin. Same three quantiles are shown. **c:** Calculation of the expected distributions of AI differences between replicates. *Top to bottom*:

– AI for each gene is calculated after pooling SNP counts from both replicates. Note that we use mean SNP coverage, so the bins contain the same genes in both replicates.
– For each coverage bin, distribution of AI values is fitted with a mixture of two symmetric beta-binomial distributions (red and blue curves in the “fit” box).
– To generate expected **Δ**AI, we perform the following steps:

– we generate a simulated sample of 5,000 genes, with the distribution of exact allelic imbalance values (ξ) according to the fitted parameters;
– from these genes, we then simulate two replicate datasets, with SNP coverage according to the bin, and sampling from binomial distribution;
– finally, we calculate the simulated **Δ**AI for each gene and find quantiles for their distribution. **d:** Plotted ratios of observed and fitted values for example quantiles, Q_65_, Q_80_, and Q_95_, calculated for coverage bins. Orange line: linear fit of the Quality Correction Constant. *Left*: Q ratios and QCC plotted against binned allelic coverage; *right*: Q ratios and QCC for each quantile.

The actual observed AI distribution has heavy tails (**Fig.2a**), suggesting that a mixture beta distribution has a better fit for allele proportions present in the sample than a single beta distribution. To gauge the experimentally observed dispersion, we performed quantile analysis of the distribution of **Δ**AI values within the coverage bins, where **Δ**AI is a difference between two replicates in AI values for a gene (**Fig.2b**).

To estimate the overdispersion, we need to normalize the observed quantile values on the expected ones. Note that genes with different AI have different impact on the overall signal dispersion (see **Suppl. Note S2**). Thus, the distribution of AI in each specific bin should be accommodated in the model.

To model the expected **Δ**AI distribution in each coverage bin and compute the corresponding quantiles, we perform the following procedure. We fit an actual distribution of AI for genes in the bin using a beta-binomial mixture model (**Fig.2c**, *top*). Using fitted parameters from that model, we then simulate two RNA-seq replicates (**Fig.2c**, *middle*). The expected distribution of **Δ**AI comes from an assumption of binomial sampling of alleles in these two simulated replicates. Finally, we calculate the quantiles for the expected **Δ**AI distribution (**Fig.2c**, *bottom*).

The ratio of observed to expected **Δ**AI quantiles appears to be a constant, with some random fluctuations (for the two replicates shown in **Fig.2d**, this ratio is 1.73 ± 0.18). This constant depends on the experiment. Poisson sampling corresponds to no overdispersion and the constant value of 1; in experimental observations we expect this value to be ≥1. We call this fitted experiment-specific quantity the Quality Correction Constant (QCC).

### Application of QCC increases concordance between replicates

To apply the experiment-specific correction, we calculate proportional test confidence intervals (CI) for allelic counts divided by QCC^2 (see Methods). Bonferroni correction for all analyzed genes is used to account for multiple hypothesis testing.

Application of this procedure to individual replicates reduced the number of genes called as imbalanced, and greatly increased the concordance between the pairs of replicate libraries (**Fig.3a**). In addition, different experiments show more similar pairwise concordance values (**Fig.3a,** top to bottom), indicating that QCC corrects for much of the experiment-specific variation. As expected, the better the agreement between technical replicates, the greater the number of imbalanced genes remaining significant (**Fig.3a**).

The observed AI values for genes with discordant balanced/imbalanced calls between two replicate libraries are now distributed closer to the boundary determined by the QCC-corrected binomial test (**Fig.3b**). This suggests that the corrected noise expectations fit the observed data better.

**Figure 3.**
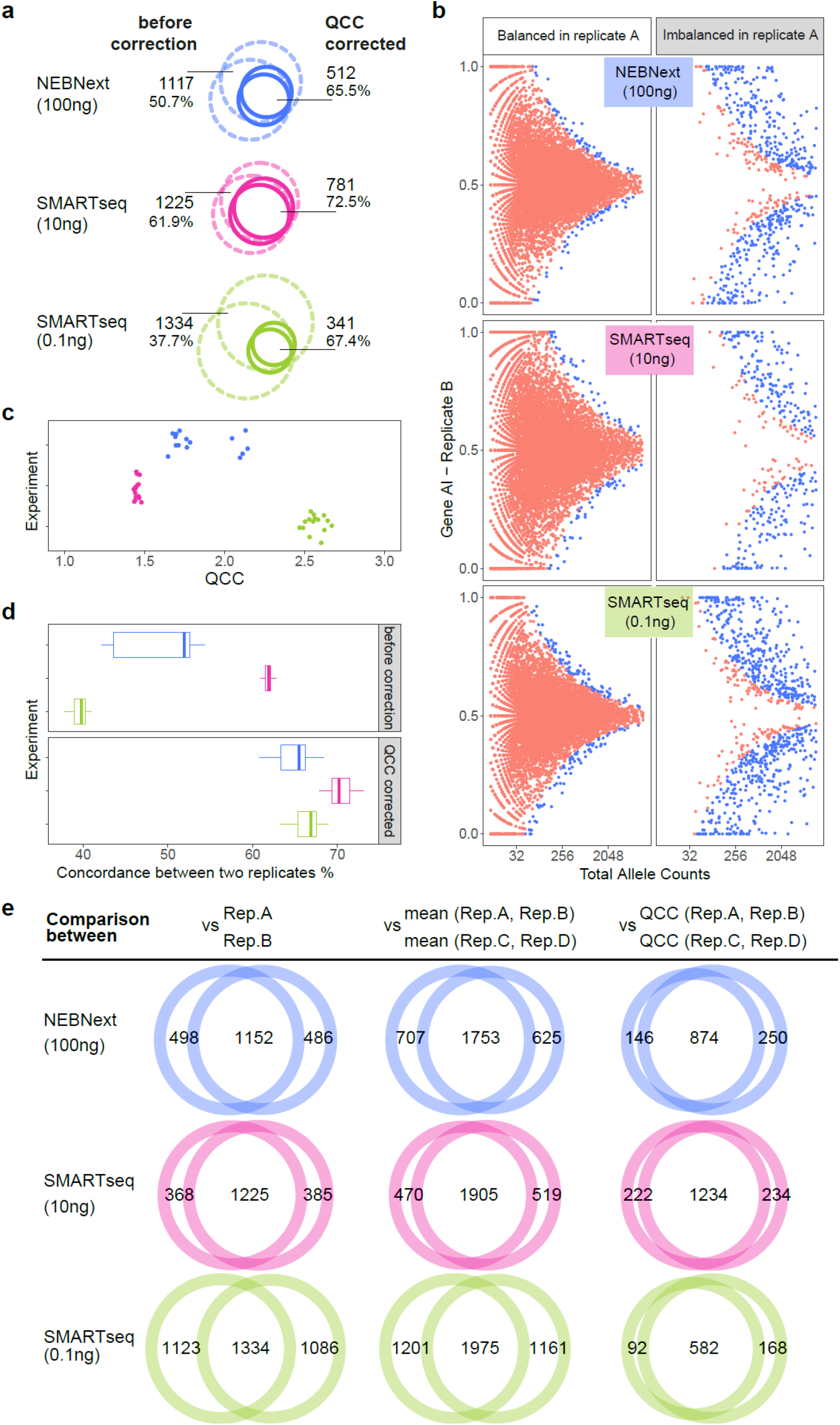
Application of QCC increases concordance between replicates and between experiments. **a:** Concordance between gene classified as allelically balanced/imbalanced between pairs of RNA-seq replicate libraries. Dotted line: using binomial test with Bonferroni correction; solid line: same replicates, using QCC-corrected binomial test. Top to bottom: NEBNext (100ng), SMARTseq (10ng), SMARTseq (0.1ng). Shown are absolute numbers of overlapping genes and % of the union constituted by the intersection. **b:** Concordance of gene classification as allelically balanced/imbalanced between pairs of RNA-seq replicate libraries after QCC correction. Genes were grouped as balanced or imbalanced based on AI values from replicate 1, and then classified as balanced (*red*) or imbalanced (*blue*) based on AI values from replicate 2. Same data used as in **Fig.1e,f,g**, re-analyzed here using QCC. **c:** QCC values calculated for all pairs of replicates within the three experiments. Color denotes experiments as in (a). **d:** Concordance for all pairs of replicates within experiments. *Top*: before QCC correction, *bottom*: with QCC correction. **e:** Concordance of balanced/imbalanced gene classification. *Left*: two replicates, using binomial test with Bonferroni correction (for each experiment, replicate #2 vs #3); *center*: pairs of pooled replicates ([#2 + #4] vs [#3 + #5]), using binomial test with Bonferroni correction; *right*: same data as in the center, with QCC correction.

To take advantage of replicate data, we can apply the correction after pooling SNP counts from the replicates, as opposed to analyzing individual replicates (as in **Fig.3a,b**). The AI point estimate thus is the weighted average between the replicates, while the coverage is the sum of the replicates’ coverage, resulting in smaller CI than in each of the replicates, and greater number of genes available for the analysis ^22,24^.

When QCC values are calculated for each pair of replicates, these values are clustered within the experiment (**Fig.3c**). This strongly suggests that it reflects an experiment-specific invariant. Note that one of 6 replicates in the NEBNext experiment appears to be an outlier, with higher QCC values when compared with all other replicates. We discuss the impact of outliers in the next section.

QCC correction led to increased concordance between replicates within each experiment and between experiments (**Fig.3d**).

Importantly, the improvement in the concordance level was due primarily to the use of the QCC, rather than the increase in coverage or pooling of the data from replicate libraries (**Fig.3e**). This reinforces the idea that the corrected, experiment-specific noise model is a better fit to the processes underlying overdispersion in RNA-seq data.

### Assessment of the correctness of the proposed approach

All the comparisons so far focused on testing the H_0_ of AI=0.5 for genes. More generally, we would like to identify genes with differential AI, and for this we need to uncover the parameters of the AI distribution in a sample. We can evaluate how close our estimates are to the true distribution by assessing the internal consistency between replicates. We thus asked if the QCC correction improves agreement between different experiments assessing the same biological sample.

To answer this question, we counted the number of genes with apparently significantly different AI (by proportional test, see Methods) when comparing pooled pairs of replicates. Since all replicates came from the same biological sample, these are false positive calls. When using only Bonferroni-corrected binomial assumptions, comparisons across experiments and even within experiments show hundreds of genes with “differential” AI (**Fig.4a**, *left*). By contrast, application of QCC removes false positives from within-experiment comparisons (**Fig.4a**, *right*). The number of false positives in across-experiment comparisons is dramatically decreased (**Fig.4a**), suggesting that QCC-corrected AI values can be used to compare AI across experiments. A nonzero number of false positives suggests the existence of systematic differences between experiments. Therefore, one should exercise caution when making comparisons across experiments

**Figure 4.**
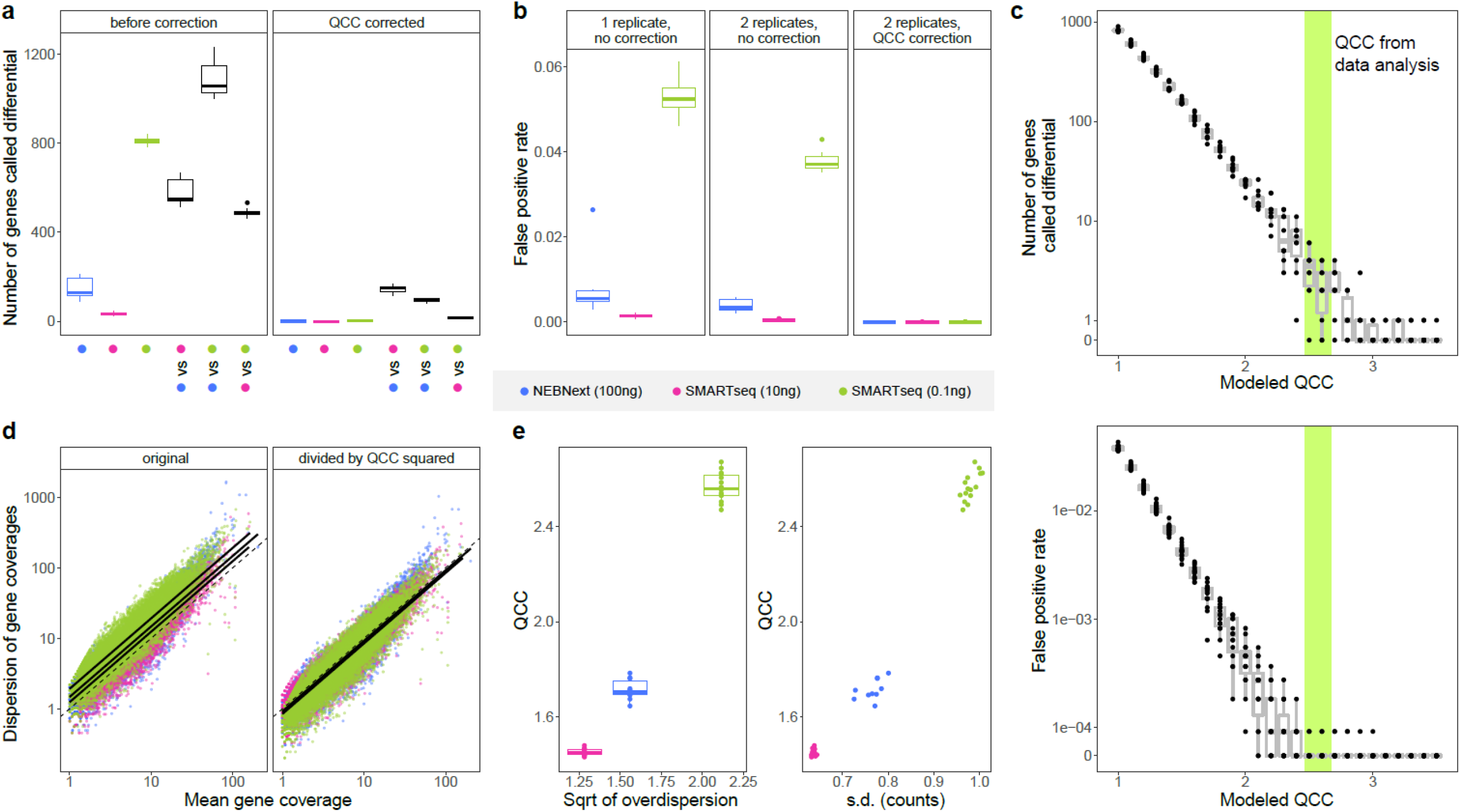
QCC enables differential AI analysis and is correlated with abundance overdispersion. **a:** Number of genes with differential expression AI in the same biological sample (i.e. false positives): before (*left*) and after QCC correction (*right*). For comparison within each of the three experiments, 15 sets of two pairs of replicates were randomly chosen within each experiment, QCC values calculated for each pair, and genes with significantly different AI in each set identified. For comparisons across experiments (*black boxplots*), the same procedure was used, except pairs were coming from distinct experiments, as noted. **b:** Impact of QCC on false positive rate. False positives are defined as genes for which the point AI estimate from 6 pooled replicates is not within CI from one replicate (*left*), two pooled replicates (*center*), and two replicates with QCC correction (*right*). **c:** QCC value is close to optimal at removing false positives. Range of calculated QCC values is contained within the colored bar. *Top* - Number of false positive genes (differential expression AI in the same biological sample) calculated for different potential values of QCC for 10 randomly selected sets of two pairs of replicates from SMARTseq (0.1ng) experiment. *Bottom* - False positive rate (as defined in b) for different possible QCC values calculated for all possible pairs of replicates in SMARTseq (0.1ng) experiment. **d:** Differences between experiments in abundance overdispersion are proportionate to QCC. *Left*: Abundance overdispersion for each experiment (color-coded as elsewhere) can be fitted as log-linear regression (*solid lines*) above expected Poisson dispersion (*dotted line*). *Right*: same after overdispersion was divided by QCC^2^. Note that the outlier replicate from NEBNext was removed for panels **d** and **e**. **e:** Correlation of QCC and abundance overdispersion. QCC values same as **Fig.3d**. Abundance overdispersion for each experiment calculated as exponent of intercept of log-linear regression (see panel d) between mean and dispersion of total counts: *Left*: for all replicates in an experiment; *Right*: for all possible pairs of replicates.

We also asked to what extent QCC correction is a better fit to the observed error distribution than the binomial assumption. We calculated the number of false positives (FP) by testing whether point AI estimates from six pooled replicates are contained within the CI of the AI estimates calculated in each of three ways. **Fig.4b** shows the FP rates for CIs obtained from one replicate under binomial assumption (*left*); pairs of replicates under binomial assumption (*middle*); and QCC-corrected from pairs of replicates (*right*). The expectation is that after the Bonferroni correction, there should be close to zero FP (see **Suppl. Note S3**). However, we observe this only after the QCC correction.

A low FP value will result from arbitrarily high QCC value, including unnecessarily high values. We asked whether the QCC value as calculated is close to optimal in that respect. **Fig.4c** shows that the computed QCC value is near the point where FP rate reaches the plateau of 0. This indicates that the corrected error model is a better fit to the underlying AI distribution in the biological sample.

It is well known that analysis of (non-allele specific) RNA abundance tends to produce data that are more variable across replicates than what is expected according to a Poisson distribution ^27–29^. We asked how this “abundance overdispersion” is related to the AI overdispersion quantified by QCC. Abundance overdispersion can be seen in all three experiments: the log-linear fit lines are above the expected Poisson dispersion (**Fig.4d**, *left*). Moreover, overdispersion was different for the three experiments. Strikingly, when dispersion values for each gene were divided by QCC^2, the regression lines for all experiments nearly coincided with each other and with the Poisson expectation (**Fig.4d**, *right*). Accordingly, abundance overdispersion was correlated with QCC values (**Fig.4e**). In simulations, QCC was also very strongly correlated with set overdispersion values (**Suppl. Fig.S2**). Based on these analyses, we hypothesize that abundance overdispersion and AI overdispersion result from largely the same processes.

While the correlation between abundance overdispersion and AI overdispersion in these examples was strong, calculation of the QCC correction is a more robust procedure that does not depend on this correlation holding for all experiments and any data processing procedure.

### Sources of AI overdispersion

To identify possible sources of AI overdispersion, we considered different stages of an RNA-seq experiment (**Suppl. Fig.S3a**): (1) steps from the biological object up to and including RNA isolation; (2) generation of sequencing library from RNA; (3) the library sequencing process itself; (4) processing of sequencing data, from read alignment to statistical analysis of allelic imbalance.

Contributions from step 1 were excluded from our experiments by design: all 18 replicate libraries were prepared from the same mouse kidney total RNA. Taking bulk aliquots of purified RNA can be considered a fair Poisson sampling process (unlike in single-cell experiments, where there are additional sources of noise such as transcription bursts ^30,31^).

Data analysis (step 4) includes multiple sub-steps, and we assess their contributions to AI overdispersion (**Suppl. Fig.S3b,c**). First, we note that these steps taken together are not a major contributing factor to variability between experiments, since input of identical data results in consistent AI and QCC values (*modulo* noise from simulation procedures).

We then asked if the tools used for allele counting and noise estimation can by themselves contribute to AI overdispersion. For an artificial example of such a contribution from the allele counting step, we applied one of the tools (such as Kallisto, RSEM and DESeq2 ^28,32–34^ that use all reads, not only reads that overlap SNPs. We expect that such read assignment should lead to a linear increase in coverage but a quadratic increase in standard deviation, and thus an increase in QCC values. Indeed, when such a tool ^34^ is applied, QCC values still cluster together, but they are systematically higher than with the ASEReadCounter* pipeline (**Suppl. Fig.S4**), which uses only the SNP-overlapping reads.

Importantly, this does not mean that ASEReadCounter* pipeline makes no contributions to AI overdispersion. This analysis would not exclude a systematic contribution to AI overdispersion which does not vary between experiments (e.g., the ASEReadCounter* pipeline counts all SNPs in a gene as independent measurements, which does not hold for SNPs found in the same read). Similarly, any allele counting procedure can have systematic biases in AI point estimates, e.g., due to reference bias in mapping (without contributing to overdispersion).

We then assessed the contribution of the process of QCC calculation to estimated AI overdispersion. On simulated total allele counts with known overdispersion, the QCC values were as expected (**Suppl. Fig.S2**; denoted as *i* in **Suppl. Fig.S3c**), suggesting that the QCC calculation step by itself makes only a very small additional contribution to noise. A related analysis starting with random binomial sampling from one replicate’s sequencing data (*ii* in **Suppl. Fig.S3c**) should show only overdispersion related to the allele counting and QCC calculation process. It yielded QCC values of 1.01-1.04 (**Suppl. Fig.S5b**), close to no overdispersion (QCC~1.0).

Note that when we randomly divide paired reads from the same run into two equal parts (cf. **Fig.1c**, *center*), these “half-replicates” are not in a binomial relationship with each other. In these comparisons (*iii* in **Suppl. Fig.S3c**), QCC values ranged from 1.45-1.48 (**Suppl. Fig.S5**), reflecting the dispersion that came in the data from one sequencing run of a single library.

Two sequencing runs with the same library (*iv* in **Suppl. Fig.S3c**), resulted in QCC values similar to half-replicates (**Suppl. Fig.S5**), suggesting that an additional sequencing run is similar to having more reads in the original run (compare *ii* and *iv* in **Suppl. Fig.S3c**). This is consistent with little, if any, contribution from the sequencing run to variation of AI overdispersion between replicates, with the caveat that we re-sequenced two libraries, and additional experiments would be needed to completely exclude sequencing process as a major source of AI overdispersion.

QCC was much larger for between-replicate comparisons than for half-replicate comparisons (*v* in **Suppl. Fig.S3c**), showing that there is additional noise coming from each replicate. Note that this underscores the point that analysis within a single replicate does not allow one to correctly account for overdispersion.

If step 1 is excluded and steps 3 and 4 are eliminated, this leaves library preparation (step 2) as the principal source of excess variability of AI estimates between replicate libraries from the same RNA. This is reinforced by the observation that technical replicate libraries have similar QCC values (see **Fig.3c**).

Generation of RNA-seq libraries involves multiple steps, from reverse transcription and cDNA fragmentation to library amplification, and these steps can substantially vary between protocols. Detailed analysis of specific protocols is outside the scope of this work. However, one common concern in deep sequencing experiments is the impact of PCR amplification artifacts ^35–37^.

To assess the impact of amplification artifacts on AI overdispersion, we compared the results of data analysis before and after removing duplicate reads. Deduplication did not reduce QCC values to ~1, and in some cases, led to increase in QCC (**Suppl. Fig.S6**) showing that other sources of noise were major contributors to AI overdispersion. Deduplication can lead to loss of large amounts of legitimate data, and may have other undesirable impacts, such as distorting signal distribution in the biological sample ^35–37^. Thus, from a practical standpoint, read deduplication has limited utility, and its impact on AI overdispersion is accounted for in the QCC analysis. Note that in paired-end RNA-seq data, the length of cDNA fragment creates unique molecular identifiers (UMI) ^38^. Thus, the results of deduplication in paired-end data (**Suppl. Fig.S6d**) suggest that the use of UMIs does not remove all AI overdispersion.

Taken together, these observations suggest that library generation is the most likely source of experiment-specific AI overdispersion and that PCR duplicates are at most partially responsible for this technical variability.

## Discussion

The replicate data we generated and the approach to testing described here can be independently used for benchmarking AI analyses. RNA-seq data from multiple replicate libraries can be used to benchmark software tools for AI estimation (see **Suppl. Fig.S4**). Conversely, QCC analysis of technical replicates of RNA-seq libraries can be used to assess the impact of different RNA-seq library preparation protocols on AI noise. It remains to be seen whether similar considerations are applicable to other sequencing assays, besides RNA sequencing.

Using these tools, we showed that variability in AI estimates between technical replicate libraries in RNA-seq experiments can be much greater than when estimated from a single replicate library (see **Fig.1c**). Importantly, this substantial AI overdispersion can vary between experiments, and thus needs to be quantified for each sample. We describe an approach that allows quantification of an experiment-specific quality correction constant (QCC) from comparison of two or more technical replicate libraries. Use of QCC results in more reproducible estimates of AI from RNA sequencing data.

Overdispersion appears uniform over all genes and thus QCC characterizes the whole experiment. QCC^2 could be thought of as a divisor for the gene allelic coverage. With observed QCC values as high as 2.67, the corrected CI are equivalent to those estimated using the (Bonferroni-corrected) beta-binomial test with fewer than 1/7^th^ of the number of sequencing reads.

It is important to note that AI overdispersion cannot be detected from a single replicate, by definition. Estimating overdispersion from one replicate requires knowing the exact underlying AI distribution to model the expected distribution and compare it with the observation. A common approximation of the underlying distribution is trimodal (AI is either exactly biallelic or completely biased towards one of the alleles; often fitted using the beta-binomial or negative binomial distribution, with the beta distribution accounting for technical noise ^17,39,40^).

However, it has long been appreciated ^41,42^ that the two alleles tend to differ in their transcriptional activity, and thus the AI of genes in a biological sample belongs to some continuous distribution on the [0, 1] interval. This makes it impossible to decompose the variation into technical and biological components using a single replicate.

With two or more technical replicates, while we still do not know the underlying AI distribution, we know it is exactly the same in the replicates, allowing us to estimate technical noise, including sampling variation and overdispersion (**Fig.2**).

Below, we describe application of QCC in two typical use cases and discuss the implications for the analysis of existing datasets lacking technical replicate libraries.

### Use case 1: One biological sample: finding genes with allelic imbalance

The simplest AI analysis of a single sample involves comparing AI of genes with a given AI value (e.g., *H*_*0*_: AI=0.5, at 0.95 confidence level with Bonferroni correction; note that this test is applied to the whole list of genes with estimated AI and CI). *H*_*0*_ for a gene is rejected if the tested AI value is outside of the CI (QCC-corrected as described in **Fig.2**).

If more than two technical replicates are available, we first calculate all pairwise QCC values for these replicates. (In a worked example in **Suppl. Note S4**, with six replicates, we obtain C_6_^2^=15 pairwise QCC values.) At this step, replicates that are outliers in quality become apparent and can be excluded from further analysis (e.g. replicate 1 of the example experiment). The mean of all the pairwise QCC values is used as the experiment-specific QCC. Point AI estimates are calculated from the pooled replicate data, and the QCC-corrected CI is used. Note that all replicates should be sampled to the same depth, determined by the replicate with the lowest number of reads; to avoid extrapolation, the safe option is to discard the extra reads from other replicate(s).

### Use case 2: Two biological samples: differential AI analysis

A more general problem is assessing if a gene has significantly different AI in two conditions. Significance is calculated using a proportional test on allelic counts corrected by respective QCC values for two samples (see Methods). Note that sampling depth used in QCC calculations does not need to be the same for all samples; thus, precomputed counts and QCC values could be directly compared.

An example comparing two clonal cell lines from 129×CastF1 mice is detailed in **Suppl. Note S5**. Note that while most comparisons discussed so far (e.g. in **Fig.1** and **Fig.3**) are based on dividing genes into discrete groups of “biased” and “unbiased”, use of corrected CI enables accurate quantification of statistically significant AI differences between samples (**Fig.4**).

### Implications for the existing data

In studies with no technical replicates available, QCC cannot be established with certainty, and thus caution should be exercised when analyzing point AI estimates. As we have shown, confidence intervals on AI estimates based on RNA-seq data depend on the experiments’ QCC. For example, we can estimate the number of genes that have AI≠0.5 in two tissue samples from a randomly chosen individual from the GTEx study ^12^. Application of the standard binomial model as in that study, with QCC=1, yields 121 such imbalanced genes for liver and 96 for lung (**Suppl. Fig.S1**). At QCC=2, these numbers would be 28 and 20 such genes, and at QCC=3, correspondingly, 19 and 11 genes.

However, very few published RNA-seq studies incorporate any technical replicates. In the Geuvadis study using human cells^22^, five samples out of 462 had technical replicates, with pairwise QCC values ranging from 1.04-1.21, lowering number of genes called imbalanced up to 1.5-fold (**Suppl. Table S2**). Considering that QCC can substantially vary even within a series of replicates (see **Fig.3** and **Fig.4**), extrapolation from these samples to the rest of those studies may not be advisable.

Biological replicates (with a single technical replicate each) are more commonly used. QCC analysis using them instead of technical replicates would give a better approximation of noise than using an implicit assumption of QCC=1. For example, in a study of allele-specific expression in mouse cells ^21^, for two samples with available biological replicates, we found QCC of 1.51 and 1.56 (**Suppl. Table S3**). Note, however, that application of QCC to biological replicates relies on the assumption that the noise between such replicates is uniformly distributed across the transcriptome, as it is for technical replicates. When this assumption is incorrect and there are actual differences between the biological replicates (e.g., a gene shifts from AI=0 to 1), calculation of QCC might lead to unpredictable errors.

## Conclusions

We described experiment-specific overdispersion that affects confidence of AI estimation in RNA-seq experiments. Consequently, it also affects confidence in differential AI analysis. This overdispersion is evident when comparing two or more technical replicate libraries, but not from the analysis of a single library, suggesting that in sequencing studies of allele-specific expression, at least two technical replicate libraries should be generated per biological sample. Our observations indicate that this experiment-specific variation mostly arises during the library construction, but amplification artifacts account for at most a fraction of it. It remains to be seen if other types of experiments besides RNA sequencing introduce similar AI overdispersion.

We describe a computational approach to account for AI overdispersion (including systematic contributions from analysis tools). This approach should increase reproducibility and functional relevance of the analyses of allele-specific expression in RNA-seq datasets.

## Methods

### 1. RNA and library preparation

Total RNA was isolated using Trizol from a freshly collected kidney tissue of an adult female mouse of 129S1 × Cast/Ei F1 background (F1 breeding was performed at the DFCI mouse facility, with parent animals obtained from the Jackson Laboratories. All animal work was performed in accordance with the institutional regulations). RNA integrity was assessed using Bioanalyzer, and it was quantified using Qubit device. Aliquots of this total RNA prep were used to prepare three sets of replicate libraries, all starting with polyA RNA isolation: 6 libraries with NEBNext kit, starting each with 100ng; 6 libraries using SMARTseq v4 kit starting with 10ng RNA; and the same, with 0.1 ng RNA. All libraries were prepared at DFCI sequencing facility according to manufacturers’ instructions. All sequencing was done on HiSeq 2500 machine at DFCI sequencing facility.

For data analysis example discussed in Use Case 2, Abelson lymphoblastoid clonal cell lines Abl.1 and Abl.2 of 129S1 × Cast/Ei F1 background ^7^ were cultured in RPMI medium (Gibco), containing 15% FBS (Sigma), 1X L-Glutamine (Gibco), 1X Penicillin/Streptomycin (Gibco) and 0.1% β-mercaptoethanol (Sigma). Total RNA was isolated extracted from cells using a magnetic bead-based protocol using Sera-Mag SpeedBeads (GE Healthcare). Isolated total RNA was DNase-treated with RQ1 DNase (Promega). RNA sequencing libraries were prepared using SMARTseq v.4 kit (Takara) starting with 10 ng total RNA for each replicate. Sequencing was performed on HiSeq4000 platform at Novogene, Inc.

Sequencing data was deposited at GEO (GSE143310).

### 2. Additional data sources

Geuvadis dataset includes RNA-seq data on LCLs established from 462 individuals from five populations ^22^.BAM files for paired-end reads (2 × 75 bp) for 5 individuals (HG00117, HG00355, NA06986, NA19095, NA20527), each with 7 replicates, were downloaded from ftp.1000genomes.ebi.ac.uk/vol1/ftp/phase3/data/.

### 3. AI estimation pipeline

AI estimation tools described here are implemented in two parts. Data processing steps from read alignment to allelic counts were based on the ASEReadCounter tool in the GATK pipeline ^16^. It was re-implemented using in part Python scripts developed by S. Castel (github.com/secastel/allelecounter), and denoted as ASEReadCounter* (github.com/gimelbrantlab/asereadcounter_star). Calculation of QCC, estimation of confidence intervals and differential AI analysis are implemented in *Qllelic* tool set (github.com/gimelbrantlab/Qllelic)

#### 3.1. Reference preparation

Two custom parental genomes (“pseudogenomes” ^43,44^; see ASEReadCounter* (github.com/gimelbrantlab/asereadcounter_star)) were used for mapping as reference. For 129S1xCast/Ei F1 cross mouse samples, alleles are determined with maternal and paternal strain genomes and strain-specific variants; for human data (Geuvadis project ^22^) phased SNP variant calls were used. Respective allelic variants from Single Nucleotide Polymorphism database 142 (dbSNP142 ^45^) or 1000 Genomes Project phase 3 structural variant call-set were inserted into the reference genome (GRCm38.86 or hs37d5, 1000 genomes, phase 2), we obtained a pair of “parental” reference genomes for further analysis (for worked example see **Suppl. Note S4**).

For each organism, we also created a vcf file with one allele considered as a reference (maternal 129S1 or first phased allele) and the other as an alternative allele. Only heterozygous sites were used in the downstream analysis.

#### 3.2. Calculation of allelic counts

##### Alignment

RNA-seq reads were aligned with STAR aligner (v.2.5.4a) ^46^ on each of two pseudogenomes, with default threshold on quality of alignment. Only uniquely aligned reads were taken into further consideration (*--outFilterMultimapNmax 1* parameter was applied).

##### Allele assignment

Reads that were present in only one of the alignments, and reads that had better alignment quality for one of the alignments, were assigned to the corresponding allele read group and marked respectively. The remaining reads (not overlapping heterozygous SNP positions) were not used downstream. This procedure is based on Python scripts by S.Castel.

##### Read deduplication

When applied, Picard (v.2.8.0; broadinstitute.github.io/picard) MarkDuplicates was used.

##### Library subsampling

To ensure that all aligned counts belong to similar distributions, BAM files corresponding to the same experiment were subsampled to the same size using a custom bash script with randomness generated using the shuf command.

##### Allelic counting for SNPs

Given a vcf file with heterozygous positions (discussed in **3.1**), coverage over each SNP was calculated using samtools mpileup (v.1.3.1) and parsed to obtain the table with allelic counts. This procedure is based on Python scripts by S.Castel.

##### Allelic counting for genes

All exons belonging to the same gene were merged into a single gene model based on GTF file (RefSeq GTF files, GRCm38.68 and GRCh37.63, were downloaded from Ensemble ftp://ftp.ensembl.org/pub/release-68/gtf/ ^47^), excluding overlapping regions that belong to multiple genes.

Phased allelic counts for all SNPs within the whole gene model were summed:

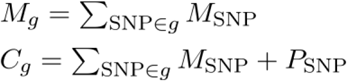

Unless specified otherwise, only genes with ≥10 total counts were used for further analysis.

##### Allelic Imbalance estimates

Estimates for AI for a gene *g* were obtained as a proportion of maternal gene counts (*M*_*g*_) to total allelic gene counts (*C*_*g*_):

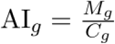

## 4. Calculation of Quality Correction Constant for 2 replicates

As gene coverage is an essential parameter of proportional beta-binomial model of allelic imbalance, we started with the standard procedure of splitting genes into bins by coverage to discretize our model.

Bin boundaries were defined as rounded up powers of the base 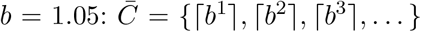. Note that QCC calculations do not strongly depend on the exact bin size, see **Suppl. Fig.S2**. Each gene *g* was assigned to a bin according to the mean of its counts 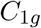 and 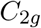 from two technical replicates:

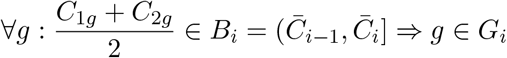

then each bin *B*_*i*_, containing set of genes *G*_*i*_, was processed separately.

### 4.1. Fitting AI distribution as beta-binomial mixture

To fit the parameters of a mixture of two proportional beta-binomial distributions, representing observed AI from the pooled replicate in each coverage bin *B*_*i*_:

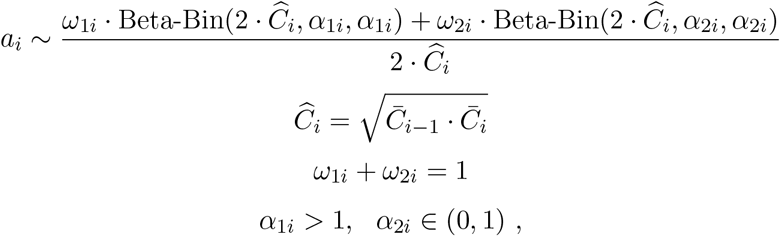

we use Expectation-maximization (EM) algorithm (see **Fig.2B**). Our fitting procedure is similar to the procedure used in the classical Gaussian mixture model ^48^.

We removed from further QCC-analysis all bins that contained less than 40 observations (genes). For fitting procedure, we also used additional threshold on the total allelic gene coverage (50 for mice and 30 for human).

Starting from initials 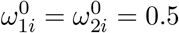, 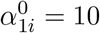, 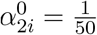, and vector of converted allelic imbalance observations 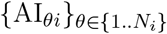, where *N*_*i*_ is number of genes in bin *B*_*i*_:

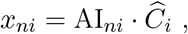

we performed iterative EM steps until the difference between parameters of the sequential steps converged (**Suppl. Fig.S7**).

#### E-step

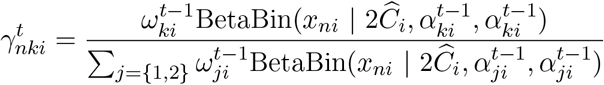

for *k* ∈ {1, 2}, *n* ∈ {1, …, *N*_*i*_} and *t* is number of EM step.

#### M-step

Since we expect 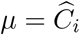 and beta-binomial distributions being symmetric:

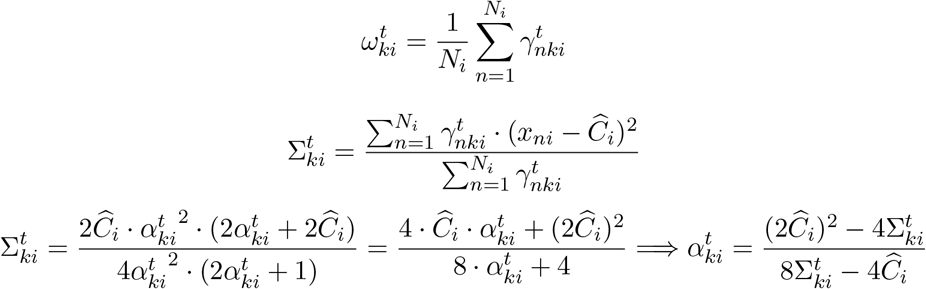

### 4.2. Simulation of a pair of replicates

Using fitted triplet of parameters {*ω*_1*i*_, *α*_1*i*_, *α*_2*i*_}, in each bin *B*_*i*_ we generated the weighted mixture of two Beta distributions probabilities {*p*_*θi*_}_*θ*∈{1..5000}_, 5000 “genes”:

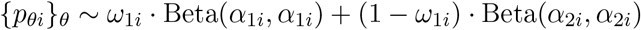

Next, for each “gene” a pair of beta-binomial distributed AIs is generated, forming two replicates.

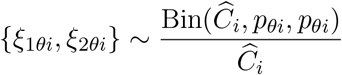

The expected AI distribution then can be obtained via subtraction: *ξ*_1*θi*_ − *ξ*_2*θi*_.

### 4.3. Quantile analysis and QCC value

To quantify the overdispersion, we performed quantile analysis between observed ΔAI distribution (**Fig.2C**) and expected ΔAI distribution (**Fig.2B**), within the coverage bins. It is a reasonable measure because differences between AI values among replicates generally tend to be symmetric on autosomes in experiments.

For each coverage bin *i* and a set of quantiles *q* ∈ {0.2, 0.35, 0.5, 0.65, 0.8, 0.9, 0.95}, the ratios of quantiles of observed ΔAI to quantiles of expected ΔAI were calculated: 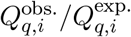.

Then the obtained ratios were linearly fitted with a constant which we call Quality Correction Constant (QCC), since it reflects the difference between observation and the binomial sampling assumption in the model (see **Fig.2D**).

## 5. More than 2 replicates in the analysis

When more than 2 replicates are used in the analysis, gene counts and AI estimates are obtained from all *M* ≥ 3 sampled replicates pooled, and the mean of all pairwise QCCs is used for correction of Confidence Intervals (CI):

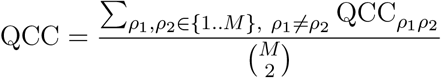

for the pair of replicates (*i, j*).

Note that before performing this step, it is useful to check if any replicates are outliers, and exclude them from further analysis.

## 6. Adjusting AI Confidence Intervals

To apply QCC and adjust CI we use `prop.test` function from R standard package `stats`, using QCC^2^ times less allelic coverage and total coverage values.

The reasoning in choosing this test is as follows: we observe that the quantiles of AI differences are QCC times wider than those from proportional binomial assumption about maternal counts distribution relative to total counts *C*_*g*_. To approximate this property for our distribution we treat gene AI observations as proportions which came from the binomial distribution for QCC^2^ times less coverage:

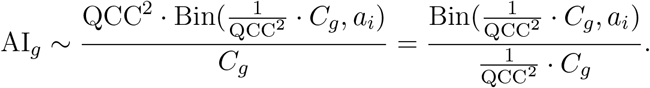

In this approximation, gene counts divided by QCC^2^ would be generally not integer, which limits the applicability of binomial test but can be addressed with proportional test which is based on Wilson score intervals.

## 7. Differential AI analysis

Accurate accounting for CIs enables differential analysis of gene AI both with point estimates and AI values from different samples.

- The difference of AI estimate from particular proportion value is considered significant if the corresponding CI interval does not cover this value.
- For identifying the differently expressed between two samples we use the the same function prop.test on the respectively corrected on the QCC values estimates as explained above.

## Supporting information

Supplementary matherials

## Acknowledgments

Supported by NIH grants HD081675 and GM114864 to AAG.

We thank Drs. M.Gelfand, A.Gusev, A.Favorov, and S.Vigneau for useful conversations and Andrew Bortvin for help with writing.

## Notes

#### Summary of Updates

Name of the tool changed to Qllelic to avoid conflicting names. Supplementary figure S4c updated to clarify the point.

https://github.com/gimelbrantlab/Qllelic

