## Supplementary matherials for "Unexpected variability of allelic imbalance estimates from RNA sequencing"

Supplementary materials for
"***Unexpected variability of allelic imbalance estimates from RNA sequencing***"


SUPPLEMENTARY TABLES 2

**Supplementary Table S1. RNA-seq datasets analyzed in this study** 2

**Supplementary Table S2. Analysis of RNA-seq from human cell lines** 3

**Supplementary Table S3. Analysis of RNA-seq from mouse neuronal progenitor cells** 4

SUPPLEMENTARY FIGURES 5

**Supplementary Figure S1. Impact of QCC value on analysis of allele-specific expression in an example GTEX dataset** 5

**Supplementary Figure S2. QCC and simulated overdispersion** 6

**Supplementary Figure S3. Principal stages of RNA-seq experiment and data analysis** 7

**Supplementary Figure S4. Sources of AI overdispersion: data processing tools** 8

**Supplementary Figure S5. Sources of AI overdispersion: impact of *in-silico* sampling and repeated sequencing runs (physical library sampling)** 9

**Supplementary Figure S6. Sources of AI overdispersion: impact of deduplication** 10

**Supplementary Figure S7. Expectation-maximization fitting of allelic imbalance Beta-Binomial mixture** 11

SUPPLEMENTARY NOTES 12

### Supplementary Table S1. RNA-seq datasets analyzed in this study

| Sample | Replicates | # fragments | Seq type | Reference |
|---|---|---|---|---|
| F1 129S1/CAST kidney, Expt. 1 (NEBnext 100ng) | 1<br>2<br>3<br>4<br>5<br>6 | 63,850,091<br>77,092,011<br>87,531,947<br>45,547,354<br>79,260,529<br>61,713,808 | SE 75 | this work,<br>GEO ID: GSE143310<br><br>Tables of allelic counts are also deposited in GEO |
| F1 129S1/CAST kidney, Expt. 2 (SMARTseq 10ng) | 1<br>2<br>3<br>4<br>5<br>6 | 85,084,237<br>73,162,672<br>65,565,213<br>74,019,699<br>73,733,135<br>90,229,986 | SE 75 | |
| F1 129S1/CAST kidney, Expt. 3 (SMARTseq 0.1 ng) | 1<br>2<br>3<br>4<br>5<br>6 | 57,996,564<br>58,816,137<br>72,585,208<br>57,195,273<br>54,874,918<br>57,807,668 | SE 75 | |
| mouse clone Abl.1 | 1<br>2 | 34,228,003<br>33,297,046 | PE 150 | |
| mouse clone Abl.2 | 1<br>2 | 33,116,795<br>33,929,680 | PE 150 | |
| mouse clone Abl.2 second round of seq | 1<br>2 | 37,381,472<br>36,658,803 | PE 150 | |
| **Data from other studies** | | | | |
| **Human RNA-seq data** | | | | |
| Geuvadis study -- see Supplementary Table S2 | | | | Lappalainen et al, (2013) Nature, 501(7468), 506-511. |
| GTEx study:<br>liver<br>lung | GTEX-11NUK-1226-SM-5P9GM<br>GTEX-11NUK-0826-SM-5HL4U | | | GTEx_Consortium. (2015). Science, 348(6235), 648-660. |
| **Mouse RNA-seq data** | | | | |
| Neuronal progenitor cells (GSE54016) -- see Supplementary Table S3 | | | | Gendrel et al, (2014) Dev Cell, 28(4), 366-380. |

**Supplementary Table S2. Analysis of RNA-seq from human cell lines**

Technical replicates were available for five samples in the Geuvadis project [Lappalainen, T., Sammeth, M., Friedländer, M. et al. Nature 501, 506–511 (2013) doi:10.1038/nature12531], with seven libraries generated from one RNA prep per each sample. Overdispersion and other AI metrics are shown for all pairwise comparisons within each replicate set (figures in bold show average and s.d.).
Allelic counts for these samples are in the Supplementary File “Allelic_counts_Geuvadis.zip”.

| sample | technical replicates | # alligned fragments in sample (PE 75) | # genes w coverage > 8 | number of genes AI != 0.5 (assuming QCC = 1) | QCC value | number of genes AI != 0.5 (with QCC) in pairwise comparisons |
|---|---|---|---|---|---|---|
| HG00117 | ERR205004<br>ERR204894<br>ERR204909<br>ERR204950<br>ERR204975<br>ERR205006<br>ERR204879 | 24,185,758<br>29,013,132<br>22,442,567<br>28,202,246<br>15,440,366<br>26,668,943<br>21,722,554 | 4609<br>4489<br>4617<br>4651<br>4351<br>4478<br>4605 | 269, 272, 272,<br>268, 275, 263,<br>285, 264, 277,<br>266, 269, 276,<br>269, 278, 272,<br>265, 273, 272,<br>280, 262, 271 | 1.104, 1.124, 1.104,<br>1.152, 1.133, 1.113,<br>1.119, 1.093, 1.108,<br>1.096, 1.14, 1.148,<br>1.163, 1.172, 1.132,<br>1.138, 1.143, 1.155,<br>1.124, 1.152, 1.162 | 225, 232, 235,<br>218, 225, 225,<br>233, 235, 228,<br>237, 215, 229,<br>216, 222, 222,<br>213, 227, 210,<br>224, 210, 213 |
| | | | **4542.9 ± 107.51** | **271.3 ± 5.85** | **1.13 ± 0.023** | **223.5 ± 8.45** |
| HG00355 | ERR204824<br>ERR204831<br>ERR204846<br>ERR204854<br>ERR204901<br>ERR204953<br>ERR204972 | 26,593,833<br>28,088,219<br>28,142,959<br>24,273,505<br>23,717,053<br>29,204,264<br>18,434,828 | 4793<br>4755<br>4737<br>4756<br>4760<br>4726<br>4503 | 212, 203, 205,<br>200, 199, 208,<br>204, 205, 194,<br>206, 205, 199,<br>195, 196, 202,<br>204, 201, 203,<br>200, 205, 198 | 1.093, 1.109, 1.082,<br>1.088, 1.106, 1.173,<br>1.126, 1.13, 1.101,<br>1.13, 1.142, 1.115,<br>1.088, 1.148, 1.169,<br>1.076, 1.126, 1.163,<br>1.09, 1.144, 1.137 | 189, 160, 179,<br>184, 170, 154,<br>167, 172, 172,<br>172, 174, 158,<br>172, 152, 147,<br>187, 160, 164,<br>171, 165, 170 |
| | | | **4718.6 ± 97.34** | **202.0 ± 4.42** | **1.12 ± 0.029** | **168.5 ± 11.04** |
| NA06986 | ERR204855<br>ERR204860<br>ERR204863<br>ERR204929<br>ERR204955<br>ERR204968<br>ERR205005 | 30,501,691<br>22,698,694<br>27,287,384<br>18,328,969<br>24,106,890<br>23,724,945<br>23,033,870 | 4654<br>4633<br>4546<br>4494<br>4616<br>4641<br>4631 | 295, 300, 304,<br>310, 308, 315,<br>301, 299, 298,<br>308, 320, 313,<br>307, 322, 316,<br>304, 322, 313,<br>316, 310, 332 | 1.125, 1.143, 1.154,<br>1.141, 1.112, 1.083,<br>1.113, 1.158, 1.165,<br>1.107, 1.108, 1.101,<br>1.129, 1.058, 1.045,<br>1.161, 1.142, 1.104,<br>1.148, 1.121, 1.098 | 248, 253, 235,<br>241, 261, 267,<br>257, 228, 228,<br>256, 270, 266,<br>253, 297, 290,<br>239, 254, 264,<br>241, 263, 280 |
| | | | **4602.1 ± 59.19** | **310.1 ± 9.35** | **1.12 ± 0.032** | **256.7 ± 18.43** |
| NA19095 | ERR204843<br>ERR204858<br>ERR204861<br>ERR204868<br>ERR204891<br>ERR204930<br>ERR205009 | 28,887,221<br>28,260,083<br>18,124,458<br>29,178,865<br>22,350,859<br>19,336,064<br>25,050333 | 5782<br>5991<br>5719<br>5901<br>5984<br>6018<br>5965 | 267, 278, 255,<br>262, 258, 274,<br>255, 262, 263,<br>256, 271, 261,<br>262, 246, 253,<br>264, 258, 271,<br>255, 279, 255 | 1.157, 1.159, 1.208,<br>1.151, 1.185, 1.192,<br>1.098, 1.094, 1.099,<br>1.111, 1.077, 1.117,<br>1.102, 1.087, 1.085,<br>1.112, 1.118, 1.122,<br>1.142, 1.103, 1.125 | 209, 211, 194,<br>203, 197, 183,<br>219, 225, 223,<br>223, 234, 215,<br>218, 213, 224,<br>219, 213, 214,<br>201, 226, 212 |
| | | | **5908.6 ± 115.22** | **262.1 ± 8.61** | **1.13 ± 0.037** | **213.1 ± 12.16** |
| NA20527 | ERR204830<br>ERR204874<br>ERR204908*<br>ERR204934<br>ERR204965<br>ERR204978<br>ERR204993 | 32,519,775<br>25,882,966<br>12,598,312<br>24,903,624<br>19,808,430<br>33,101,465<br>27,981,881 | 4766<br>4849<br>4728<br>4825<br>4862<br>4836<br>4855 | 213, 211, 209,<br>220, 204, 206,<br>208, 212, 211,<br>204, 208, 213,<br>222, 210, 209,<br>205, 207, 219,<br>205, 212, 201 | 1.097, 1.063, 1.077,<br>1.118, 1.15, 1.068,<br>1.11, 1.11, 1.141,<br>1.079, 1.112, 1.123,<br>1.176, 1.132, 1.133,<br>1.12, 1.188, 1.107,<br>1.177, 1.107, 1.095 | 183, 189, 184,<br>177, 166, 186,<br>178, 187, 167,<br>190, 183, 187,<br>168, 173, 176,<br>179, 158, 186,<br>153, 178, 177 |
| | | | **4817.29 ± 50.73** | **210.0 ± 5.44** | **1.12 ± 0.034** | **177.4 ± 10.14** |

* - this sample had the fewest reads; to ensure uniform analysis, all the sampling was performed to this depth.

**Supplementary Table S3. Analysis of RNA-seq from mouse neuronal progenitor cells**

Two biological replicates were available for two samples [Gendrel, A.V. et al. Dev Cell. 2014 Feb 24;28(4):366-80 doi: 10.1016/j.devcel.2014.01.016]. Overdispersion and other AI metrics are shown for a replicate pair.
Allelic counts for these samples are in the Supplementary File “Allelic_counts_NPC.zip”.

| sample | biological replicates | # alligned fragments in sample (PE 100) | # genes w coverage > 8 | number of genes AI != 0.5 (with QCC = 1) | QCC value | number of genes AI != 0.5 (with QCC) in pairwise comparisons |
|---|---|---|---|---|---|---|
| SRS529152<br>SRS529162 | SRR1106776<br>SRR1106785 | 79,524,208<br>169,065,784 | 12531<br>12661 | 3338 | 1.51 | 1958 |
| SRS529159<br>SRS529163 | SRR1106781<br>SRR1106786* | 75,319,044<br>26,302,221 | 12383<br>12373 | 3104 | 1.56 | 1665 |

* - this sample had the fewest reads; to ensure uniform analysis, all the sampling was performed to this depth.

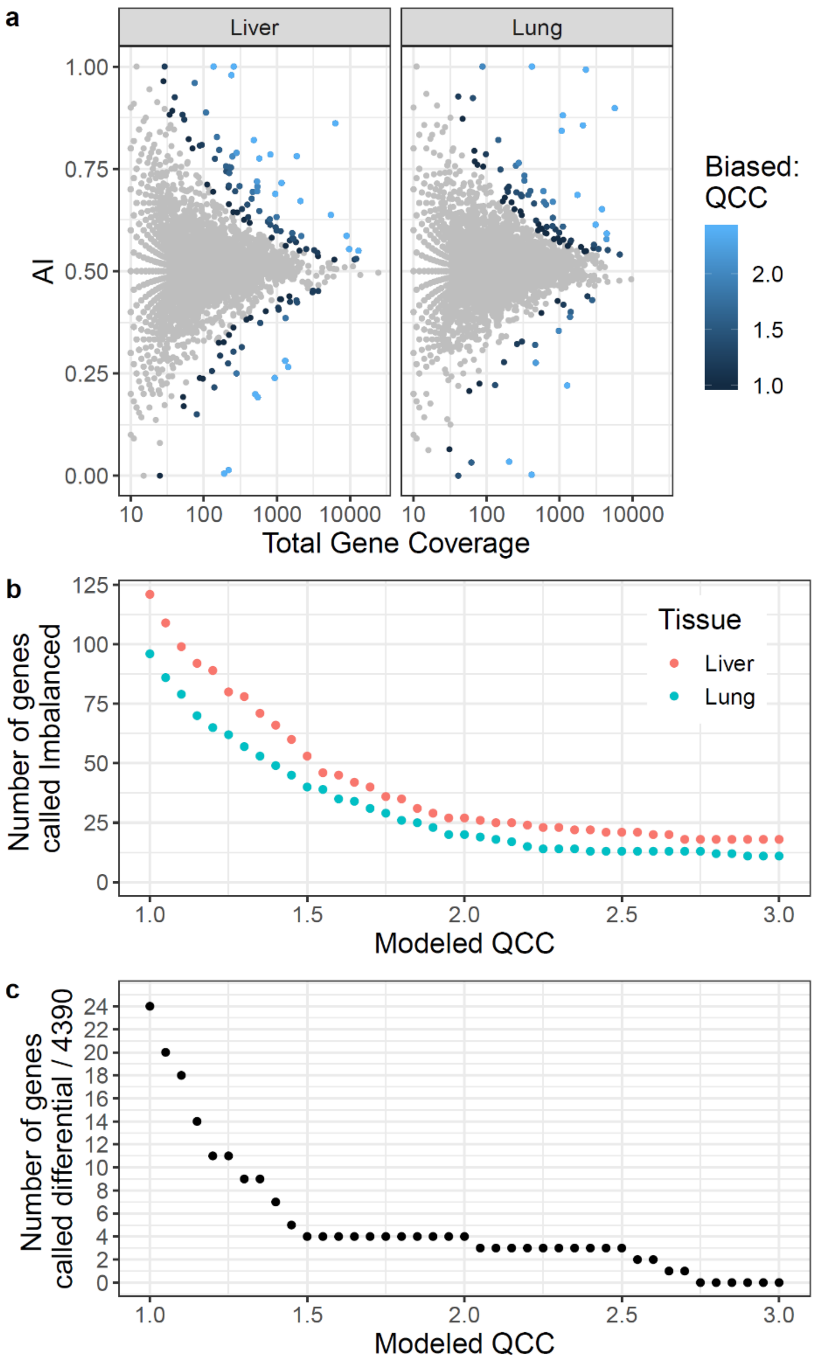


**Supplementary Figure S1. Impact of QCC value on analysis of allele-specific expression in an example GTEX dataset**

To illustrate the impact of overdispersion on the analysis of expression AI in a dataset with no replicates available, we assessed RNA-seq data for two human tissues, liver and lung, from the same arbitrarily selected individual (GTEX-11NUK) from the GTEx project (no replicates available). Allelic counts for these samples are in the Supplementary File “Allelic_counts_GTEx_11NUK.zip”.

**a:** Distribution of significantly biased genes (testing $H_0$ of AI=0.5) at different assessed QCC values from 1.0 (no overdispersion) to 3.0.

**b:** Same analysis as A, showing the number of imbalanced genes for the two tissues.

**c:** Differential analysis of AI expression between the two tissues, when corrected using different QCC values (equal for both samples).

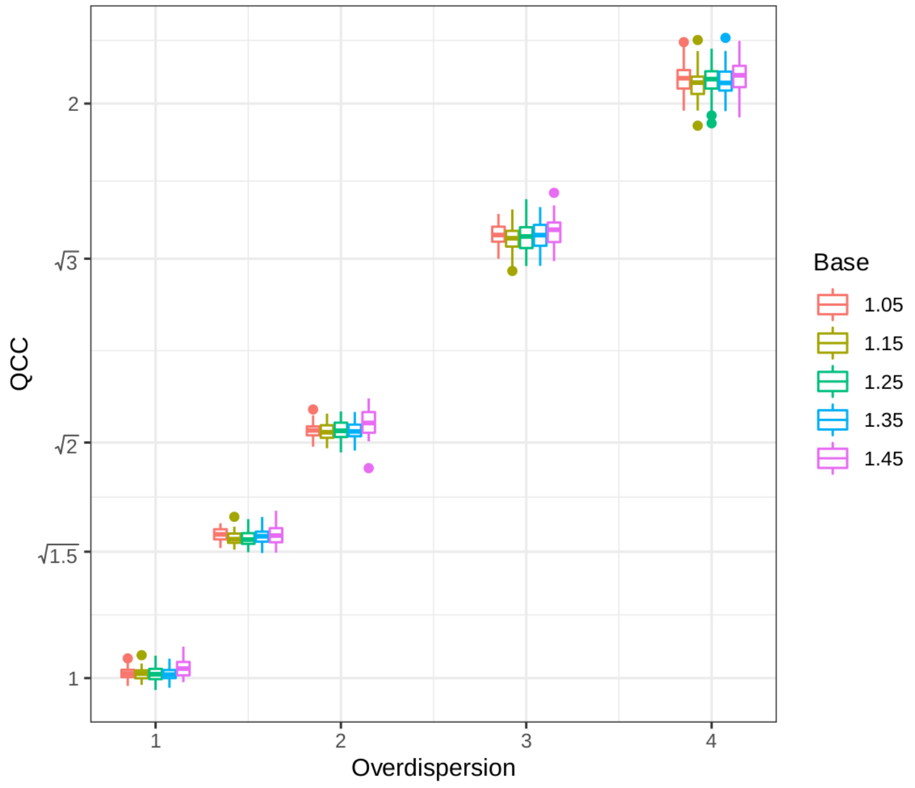


**Supplementary Figure S2. QCC and simulated overdispersion**

Gene coverage bin size doesn't substantially change QCC estimate. QCC values were calculated for simulated data with preset overdispersion values (horizontal axis), for a range of coverage bin sizes (color-coded by exponential bin base as noted). Note that calculated overdispersion values (QCC) nearly perfectly correlated with overdispersion values set in the simulation.

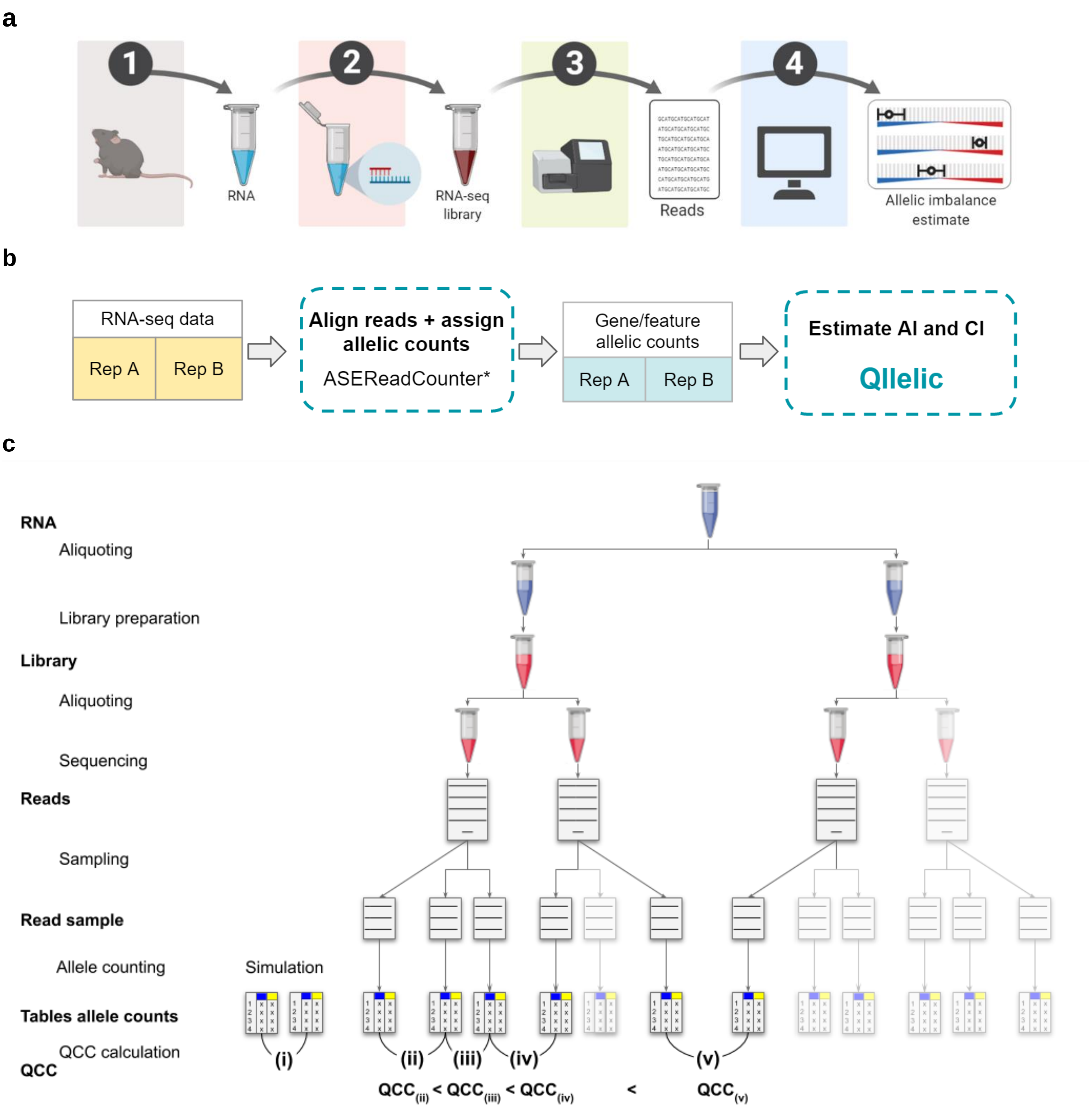


### Supplementary Figure S3. Principal stages of RNA-seq experiment and data analysis

**a:** Steps of RNA-seq experiment. 1 – RNA isolation; 2 – Library preparation; 3 – Sequencing, resulting in “reads” (which can be pairs in PE sequencing). Separate sequencing runs of the same library result in “physical subsampling” of the library; 4 – Data analysis

**b:** Overview of data analysis with ASEReadCounter* and Qllelic. Note that other tools could be used for allele counting instead of ASEReadCounter* (e.g., see **Suppl. Fig.S4**).

**c:** How datasets were prepared for analysis of source of overdispersion in data processing. Comparisons (i) – (v) are described in detail in Discussion. For specific QCC values for these comparisons, see: (i) – **Suppl. Fig.S2**; (ii),(iii),(v) – **Suppl.Fig.S5b**; (iii),(iv),(v) - **Suppl.Fig.S5c.**

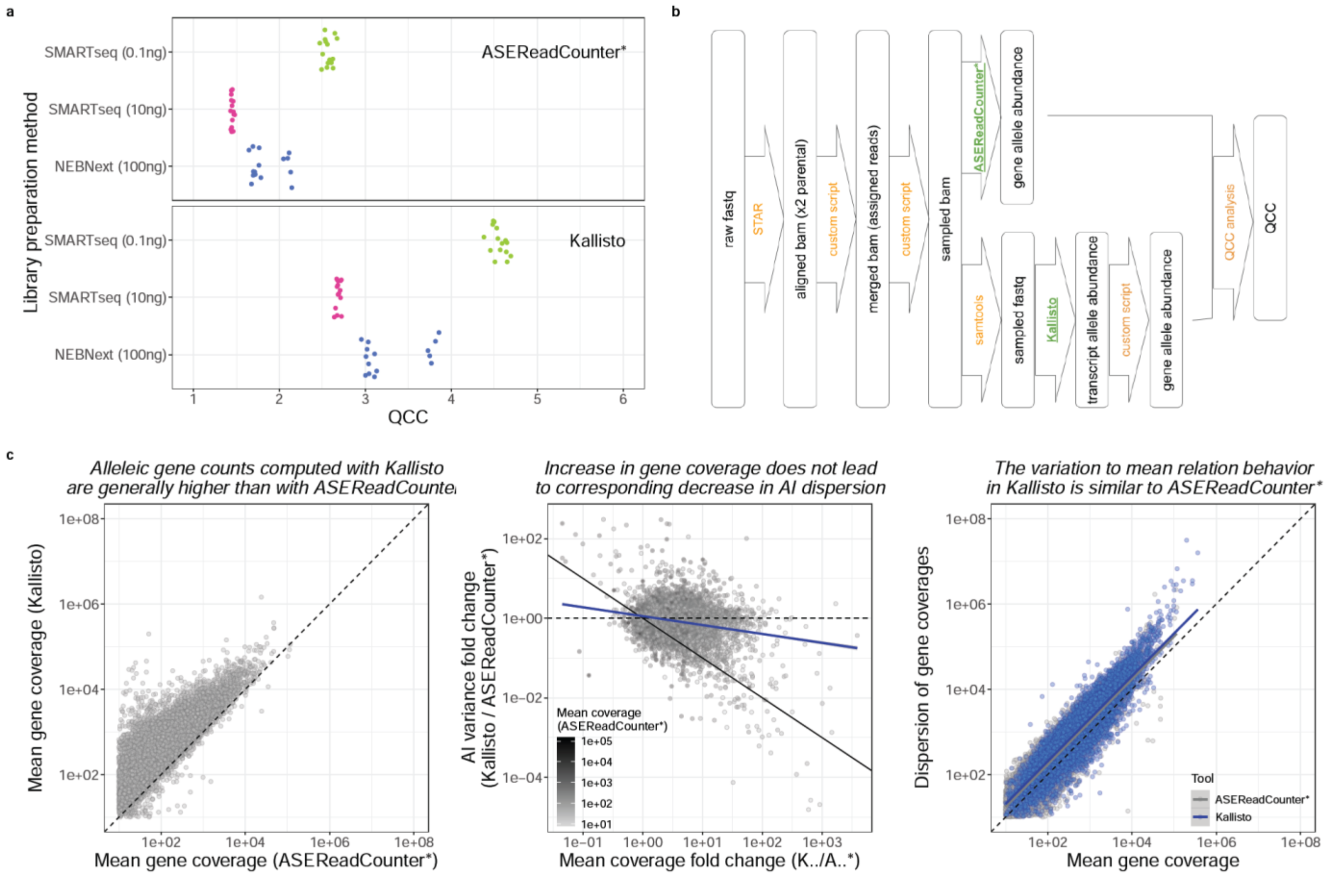


### Supplementary Figure S4. Sources of AI overdispersion: data processing tools

**a:** QCC values calculated for allelic counts computed from the same set of aligned reads by the default counter (ASEReadCounter*) and another tool (Kallisto).

**b:** Outline of data processing with ASEReadCounter* and Kallisto.

**c:** Source of elevated overdispersion when using Kallisto for allelic coverage counting.
*Left* - Kallisto pipeline yields higher allelic counts (due to assignment of non-SNP-covering reads to allelic counts).
*Center* - Despite higher allelic counts in Kallisto, no great decrease in AI variance (black solid line - expected relationship between fold change in AI variance and fold change in mean gene coverage change; dashed line - expected if AI variance were independent of mean coverage; blue line - linear fit of observed relationship).
*Right* - Relationship between gene coverage and AI dispersion is similar for Kallisto (blue) and ASEReadCounter* (grey). Also compare to **Fig.4D**.

For this figure, analyses performed on 30M reads sampled from each of six replicates in the SMARTseq (10 ng) dataset.

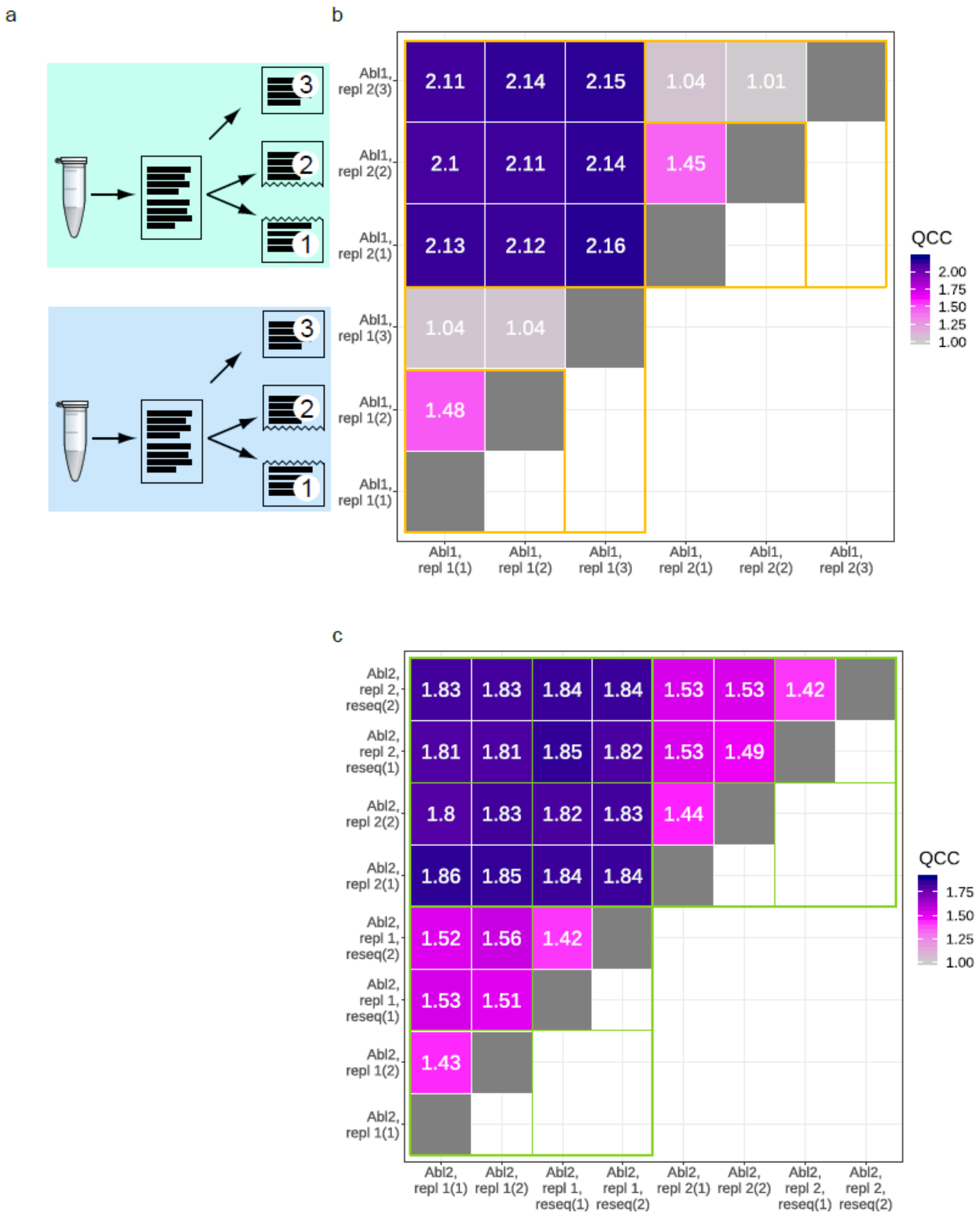


**Supplementary Figure S5. Sources of AI overdispersion: impact of *in-silico* sampling and repeated sequencing runs (physical library sampling)**

**a:** Schematics of the sampling. “1” and “2” are two non-intersecting subsets of reads [see definition in **Fig.S3**] from the same sequencing run of a particular replicate after alignment and assignment to parental genomes (paired reads were kept together). “3” is an independent sampling (of the same size as “1” and “2”) of the same pool of reads. It is in binomial relationship with each of “1” and “2”, e.g., it may overlap with these subsets.

**b:** For within-replicate comparison, *in-silico* sampling without return has greater contribution to AI overdispersion than binomial sampling. Pairwise QCC analysis performed on Abl.1 data, replicates 1 and 2. Subsets “1” and “2” are non-intersecting subsets (15,136,606 fragments each) of a given library sequencing run; “3” random subset of 15,136,606 fragments from the same sequencing run. Orange boxes highlight comparisons between “1” and “2” and between them and “3”.

**c:** Repeated sequencing runs of the same library (“physical sampling”) has small impact on AI overdispersion. Pairwise QCC analysis on Abl.2 data, replicates 1 and 2. Resequencing as annotated. Green boxes highlight comparisons within and between resequencing runs.

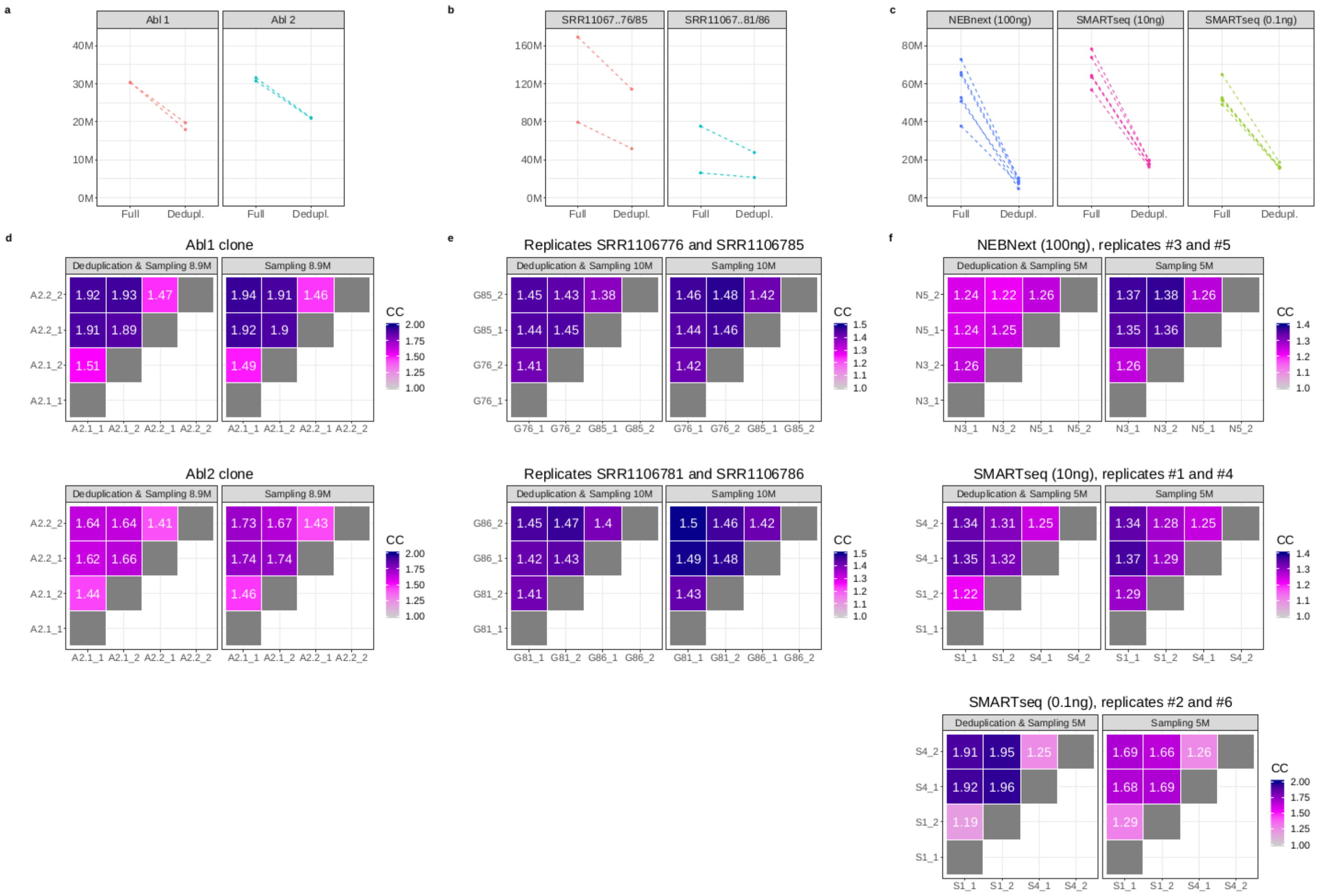


### Supplementary Figure S6. Sources of AI overdispersion: impact of deduplication

**a, b, c:** Number of fragments remaining after deduplication (using Picard MarkDuplicates).
a - PE150 data from Abl.1 and Abl.2
b - PE75 data from NPC cells (Gendrel dataset)
c - SE75 data for the kidney RNA-seq data in Experiments 1-3.

**d, e, f:** QCC after deduplication is still higher for comparing two replicates than comparing halves of one replicate.
d - QCC before and after deduplication: Abl.1 and Abl.2 data
e - QCC before and after deduplication: NPC cells (Gendrel dataset)
f - QCC before and after deduplication for randomly chosen pairs of replicates of kidney RNA-seq data. Note that QCC can become higher after deduplication.

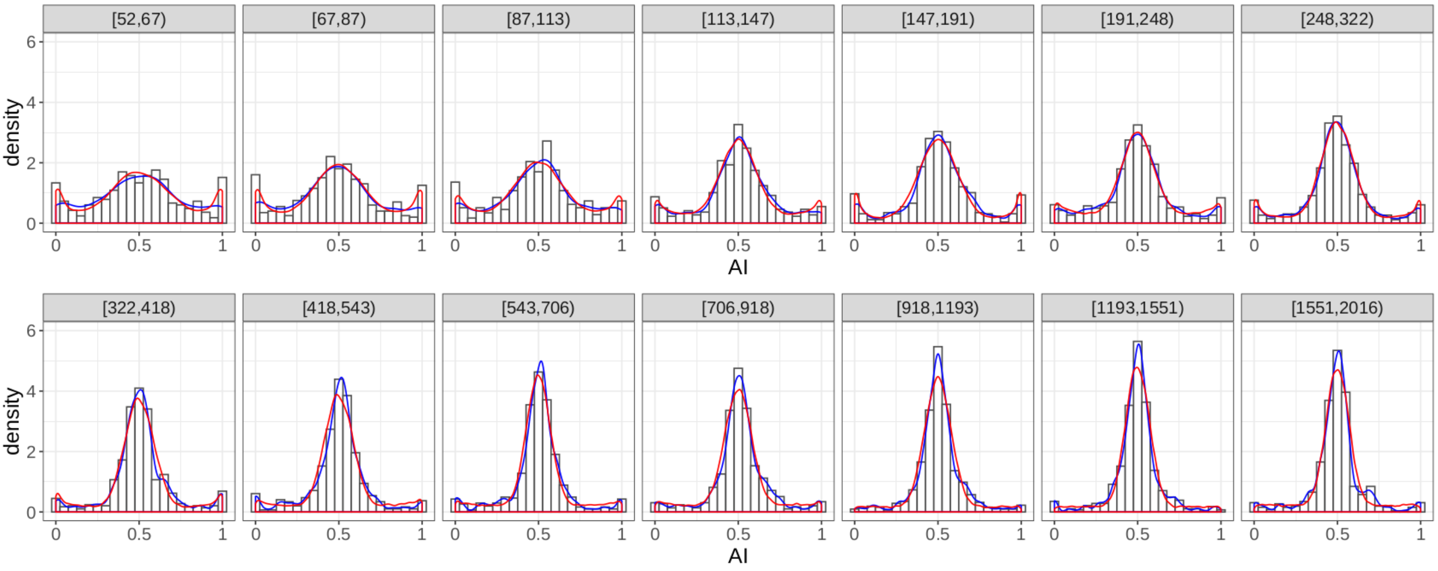


**Supplementary Figure S7. Expectation-maximization fitting of allelic imbalance Beta-Binomial mixture**

Shown are gene coverage bins (as annotated in each box). The *histogram* is the observed AI distribution in the actual sample in each bin; *blue line* is smoothed density plot of the histogram; *red line* shows fitted distribution. EM algorithm applied to non-deduplicated RNA-seq data for Abl.1 clone, sampled to 30,273,212 PE150 reads in each technical replicate; base of exponential binning is 1.3.

### Supplementary Note S1: Accounting for overdispersion leads to the expected bimodal distribution of AI values in discordant AI calls.

With what probability the binomial-like test (**Fig.SN1a**) would show different results on two maternal counts observations $M_1$ and $M_2$ for a given underlying proportion $a$ (**Fig.SN1b**)?

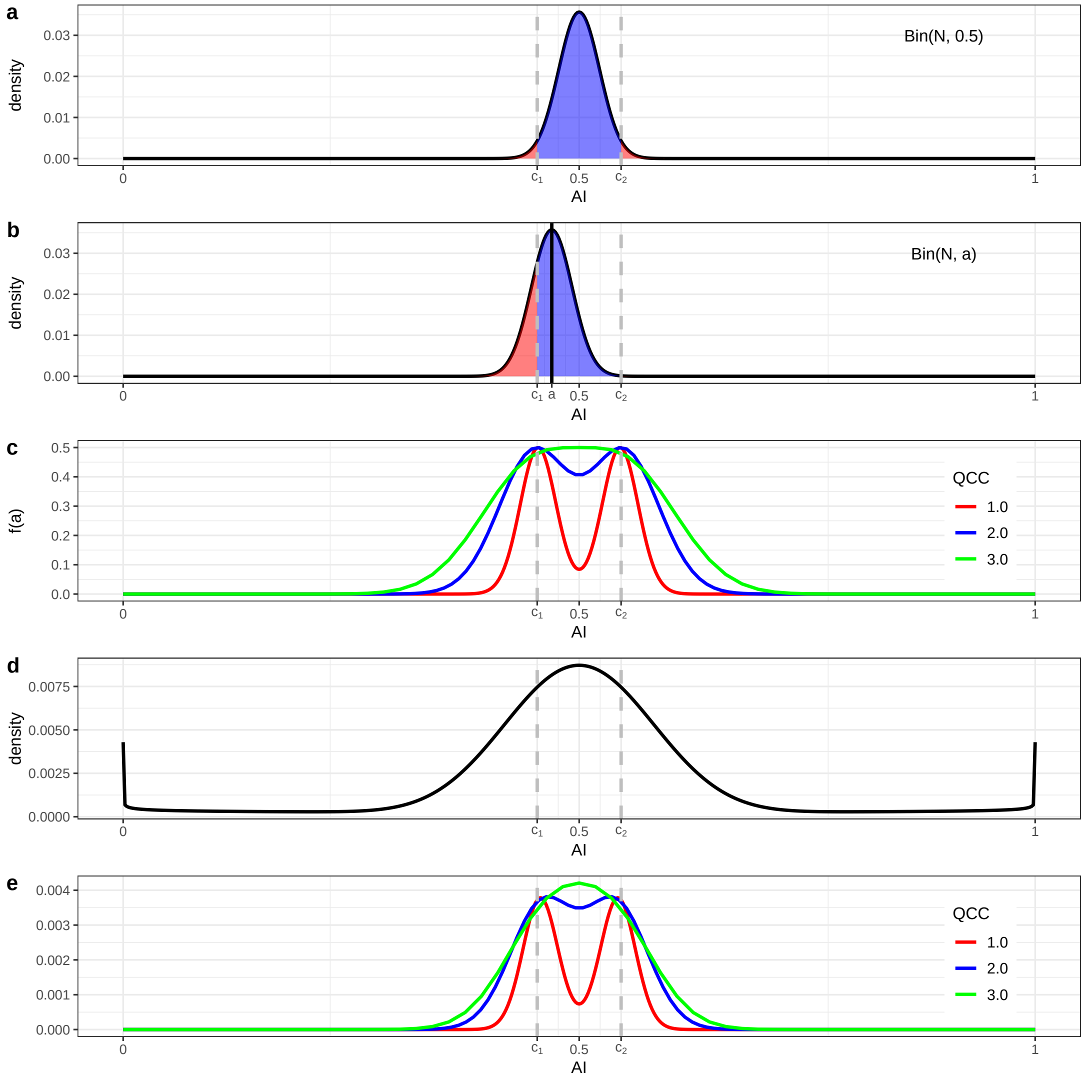


Figure. SN 1: **(a)** Binomial distribution and boundaries of binomial test with 0.5, for given $N = 500$ and level of confidence 0.95;
**(b)** Binomial distribution of observed AI for underlying imbalance $a = 0.47$, colored according to binomial test result;
**(c)** Probability to receive discordant test results for 2 observations for different $QCC$ (recall that $QCC = 1$ corresponds to binomial distribution and thus represents the case when the test fits the data);
**(d)** Sample distribution of underlying AI;
**(e)** Distribution of underlying AI of genes that may be marked differently when doing binomial test for 2 technical replicates (for $QCC = 1$ we see that it is bimodal and the peaks are near binomial test boundaries).

Let us consider the function of that probability for particular gene coverage N and respective boundaries $C_1$ and $C_2$ of the binomial test $\mathrm{BT}_{QCC}$ with $H_0 : p = 0.5$, on counts respectivelly corrected on QCC (**Fig.SN1c**). Then probability of discordant True/False results of $\mathrm{BT}_{QCC}$ on 2 technical replicates for underlying proportions $a \in (0, 1)$ is:

$$f_{N,QCC}(a) = P(\mathrm{BT}_{QCC}(M_1) \neq \mathrm{BT}_{QCC}(M_2)) =$$

$$= 2 \cdot (P(M \leq C_1|a) \cdot P(M \in (C_1, C_2)|a) + P(M \geq C_2|a) \cdot P(M \in (C_1, C_2)|a)) =$$

$$= 2 \cdot P(M \in (C_1, C_2)|a) \cdot (P(M \leq C_1|a) + P(M \geq C_2|a)) =$$

$$= 2 \cdot \int_{C_1}^{C_2} \mathrm{Bin}_{QCC}(x; N, a)dx \cdot (\int_{0}^{C_1} \mathrm{Bin}_{QCC}(x; N, a)dx + \int_{C_2}^{N} \mathrm{Bin}_{QCC}(x; N, a)dx)$$

Given discretized distribution $U_N(a)$ of underlying AI (**Fig.SN1d**), we may obtain the distribution of AI of genes with discordant results of $\mathrm{BT}_{QCC}$ on 2 technical replicates, as a product of $U_N(a)$ and $f_{N,QCC}(a)$ (**Fig.SN1e**).

Note that if the test uses the distribution which is relevant to the data, the resulting distribution of AI of genes with discordant results will be distributed around test boundaries. By contrast, not accounting for overdispersion tends to much wider, unimodal distribution (which is what we observe in **Fig.1e-g**).

### Supplementary Note S2: Genes with different underlying AI have different impact on the overall signal variance.

Let us consider genes in particular coverage bin.

If AI values for 2 given replicates, $x_1 = \{x_{1i}\}$ and $x_2 = \{x_{2i}\}$, belong to the same binomial-like distribution and corresponding underlying allelic proportions $a = \{a_i\}$ belong to any symmetric distribution, then since $var(x_{1i} - x_{2i}) = var(x_{1i}) + var(x_{2i})$ we expect that $var(x_{1i} - x_{2i}) \sim a_i(1 - a_i)$ for any gene $i$.

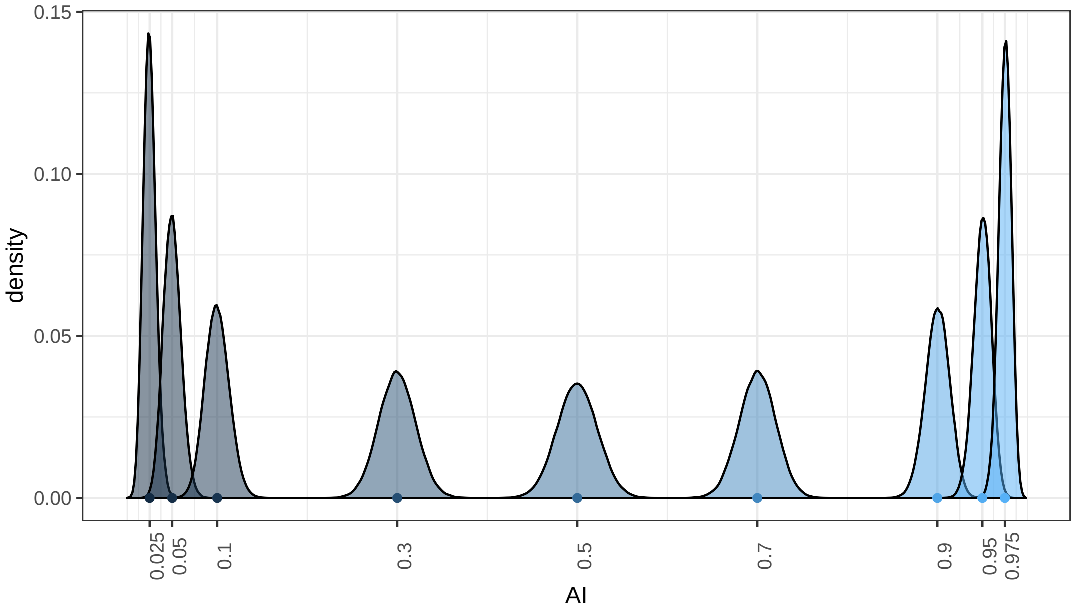


Figure. SN 2: Variance of AI observations differs along the interlval of underlying AI (gene coverage 500).

And $a_i(1 - a_i)$ reaches its maximum at 0.5 and minimum at 0 and 1 (**Fig.SN2**).

For $X = \{x_{1i} - x_{2i}\}_i$ then $var(X) = E((X - \mu)^2) = E((X - 0)^2) = \frac{1}{n}\sum_{i=1}^{n}(x_{1i} - x_{2i})^2$, which depend on the distribution of underlying allelic proportions $a = \{a_i\}$.

And since underlying AI belong to some complex distribution, and genes with different underlying AI have different impact on the overall signal variance, the AI distribution should be taken into account when making any conclusions about distribution of differences between AI observations in technical replicates.

Thus we conclude that it is necessary to account for the distribution of underlying allelic imbalances when assessing AI differences between technical replicates, while assumption of a simple distribution (e.g. tri-modal) is insufficient for this purpose. We account for the observed distribution in the beta-binomial model (see **Fig.2C**).

### Supplementary Note S3: We expect nearly zero genes with false positive AI, when we estimate AI and CI from two replicates and then calculate AI from six replicates.

Here, we define "false positive" as the event when the point estimate of AI for a gene from six replicates is outside of the CI for the same gene based on two replicates out of six (after Bonferroni correction).

For particular gene and respective underlying AI $p$, maternal counts in $n$ replicates are expected to follow the same distribution $m_{i\in\{1..n\}} \sim \text{Bin}(C,p)$ (we consider here the case of QCC $= 1$, but it can be clearly generalized). Then for $\text{CI}(\text{AI}_2 = \frac{m_1+m_2}{2\cdot C})$ computed on the first 2 replicates, the probability of underlying AI $p$ to miss $\text{CI}(\text{AI}_2)$ (which is determined by confidence level by design, for example, 0.05) is less than probability of AI estimate on $k$ replicates that are different from the first 2 replicates $(\frac{\sum_3^{k+2} m_i}{k\cdot C} \sim \frac{\text{Bin}(k\cdot C,p)}{k\cdot C})$, but is greater than probability of $\text{CI}(\text{AI}_2)$ to be missed by AI estimate computed on $k$ replicates that include the first 2 replicates $(\frac{\sum_1^k m_i}{k\cdot C} \sim \frac{\text{Bin}(k\cdot C,p)}{k\cdot C})$, and not independent with $\frac{m_1+m_2}{2\cdot C})$.

In our case:

$$P\left(\frac{\sum_1^6 m_i}{6\cdot C} \notin \text{CI}\left(\frac{m_1+m_2}{2\cdot C}\right)\right) \leq P\left(p \notin \text{CI}\left(\frac{m_1+m_2}{2\cdot C}\right)\right) \leq P\left(\frac{\sum_3^8 m_i}{6\cdot C} \notin \text{CI}\left(\frac{m_1+m_2}{2\cdot C}\right)\right)$$

Simulations supporting this idea are provided in **Fig.SN3** and **Fig.SN4**.

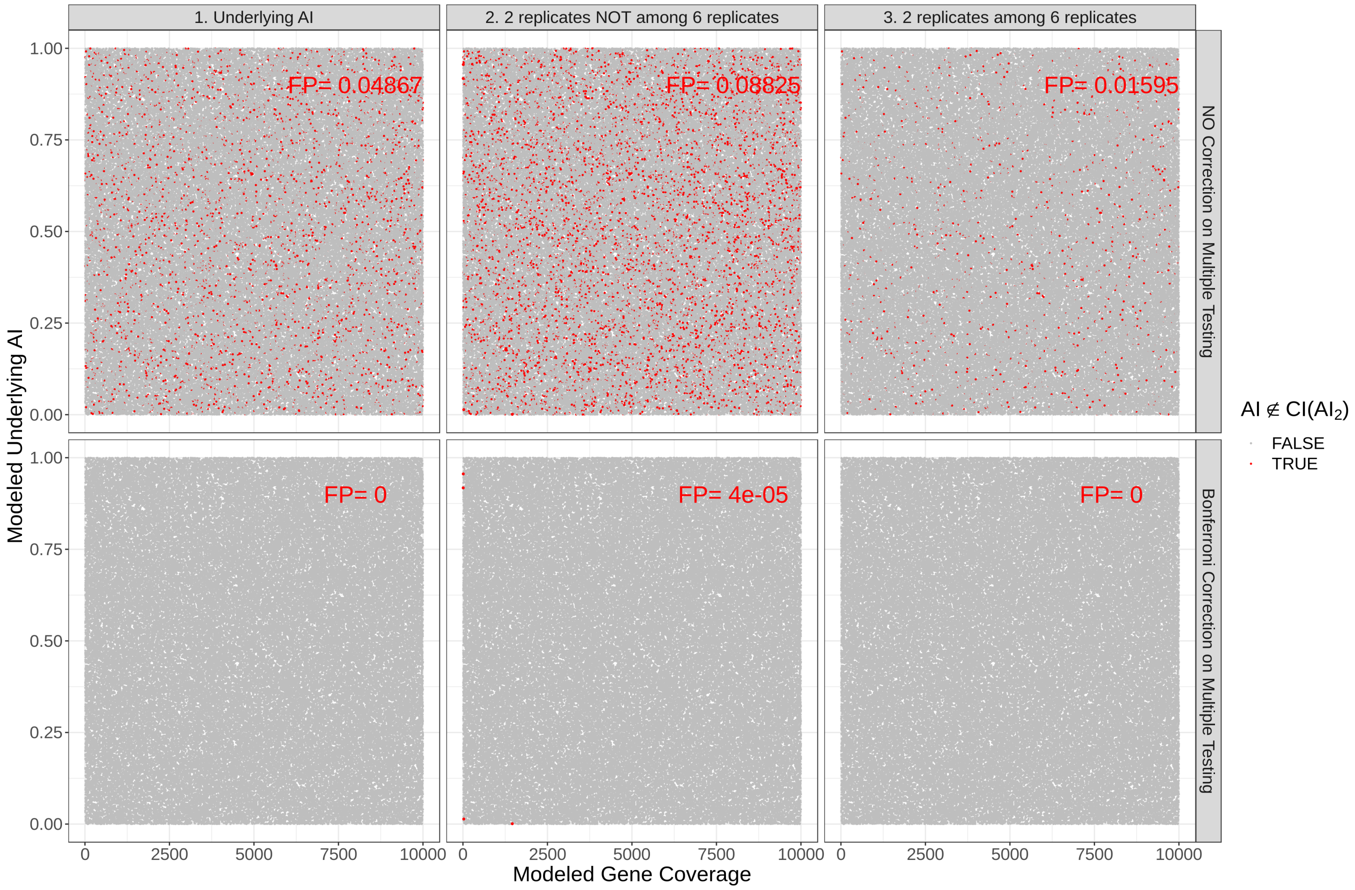


Figure. SN 3: Number of false positives for modeled 100,000 genes with different coverage and underlying AI. The point used to test if it lies within $CI(AI_2)$ calculated on 2 randomly selected replicates, left to right: underlying AI, AI estimate from a set of 6 replicates that are different from initial 2 replicates, AI estimate from a set of 6 replicates which contain initial 2 replicates. The false positive rate is provided before and after correction on multiple testing.

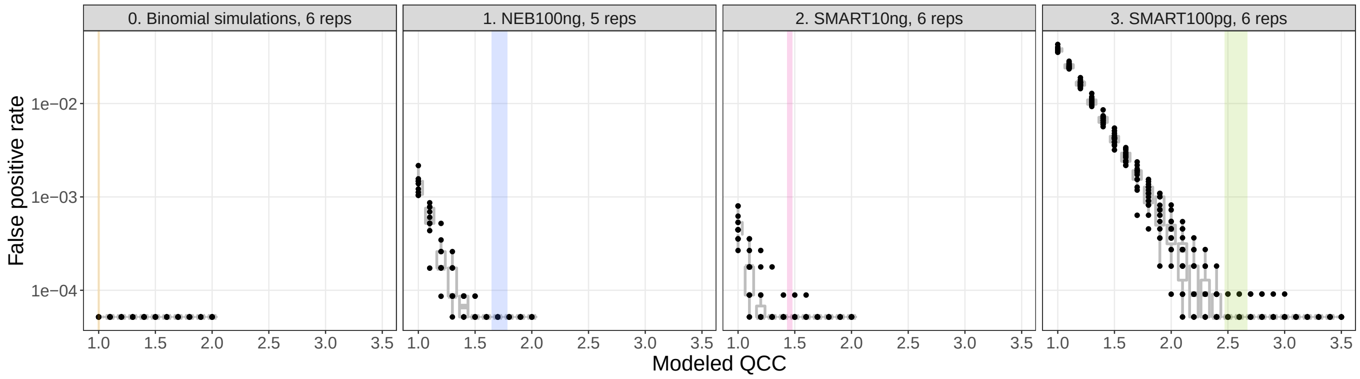


Figure. SN 4: False positive rate for binomial simulations ($QCC = 1$) and different experiments for different modeled QCC. The range of real QCC is colored.

**Supplementary Note S4: Worked example for QCC calculation, starting from fastq**
fastq -> counts: https://github.com/gimelbrantlab/ASEReadCounter_star/wiki/2.-Allelic-Counts-Table-Creation
counts -> QCC: https://github.com/gimelbrantlab/Qllelic/wiki/Use-case-1:-One-biological-sample

**Supplementary Note S5: Worked example of AI differential analysis for two samples**
https://github.com/gimelbrantlab/Qllelic/wiki/Use-case-2:-Differential-AI-analysis
